# Extensive profiling of transcription factors in postmortem brains defines genomic occupancy in disease-relevant cell types and links TF activities to neuropsychiatric disorders

**DOI:** 10.1101/2023.06.21.545934

**Authors:** Jacob M. Loupe, Ashlyn G. Anderson, Lindsay F. Rizzardi, Ivan Rodriguez-Nunez, Belle Moyers, Katie Trausch-Lowther, Rashmi Jain, William E. Bunney, Blynn G. Bunney, Preston Cartagena, Adolfo Sequeira, Stanley J. Watson, Huda Akil, Gregory M. Cooper, Richard M. Myers

**Affiliations:** HudsonAlpha Institute for Biotechnology, Huntsville, AL, USA; Department of Psychiatry and Human Behavior, University of California, Irvine, CA, USA; The Michigan Neuroscience Institute, University of Michigan, Ann Arbor MI, USA

**Keywords:** Transcription Factors, ChIP-seq, Neuropsychiatric Disease, Brain, *cis*-regulatory elements

## Abstract

Transcription factors (TFs) orchestrate gene expression programs crucial for cell physiology, but our knowledge of their function in the brain is limited. Using bulk tissues and sorted nuclei from multiple human post-mortem brain regions, we generated a multi-omic resource (1121 total experiments) that includes binding maps for more than 100 TFs. We demonstrate improved measurements of TF activity, including motif recognition and gene expression modeling, upon identification and removal of regions of high TF occupancy. Further, we find that predictive TF binding models demonstrate a bias for these high occupancy sites. Neuronal TFs SATB2 and TBR1 bind unique regions depleted for such sites and promote neuronal gene expression. Several TFs, including TBR1 and PKNOX1, are enriched for risk variants associated with neuropsychiatric disorders, predominantly in neurons. These data are a powerful resource for future studies seeking to understand the role of TFs in epigenetic regulation in the human brain.

## Introduction

Transcription factors (TFs) are a major class of DNA-associated proteins that also include transcriptional cofactors and chromatin remodelers^1, 2^. These proteins play critical roles in every biological process, including development, cell fate determination, and physiological responses^3–6^. They carry out these functions primarily by localizing to distinct genomic regions, typically referred to as *cis*-regulatory elements (CREs), and regulate gene transcription. Disruption or alteration of TF-mediated effects can increase disease risk^7, 8^, making the study of TF functions a critical component of understanding health and disease. Several TFs with critical roles in normal brain development are involved with neuropsychiatric disorders, such as SP4^9^, EGR1^10^ NR3C1^11^, TCF4^12^, and additional TFs are implicated through recent GWASs^13–16^. CREs in brain tissues are enriched for GWAS hits for multiple neurological disorders, suggesting they share genetic risk factors that act through altered gene regulation^17–19^. Additional analysis has shown that multiple psychiatric disorders associate with the same loci^20^. Because of their prominent role in gene regulation, understanding TF functions is a critical component of understanding brain health and disease. Recent work by the PsychENCODE^21^ and CommonMind^22^ consortia has generated reference maps for promoter (H3K4me3) and enhancer (H3K27ac) associated histone modifications from cortical tissues of donors. These data show enrichment for neuronal development and neuropsychiatric risk variants in regions with these chromatin marks^23, 24^. While the ENCODE Consortium^25^ has generated thousands of occupancy maps for human TFs, these are mostly derived from a small set of cell lines that are distinct from the primary tissues relevant for many diseases. Few TFs have been directly studied in human brain tissues, particularly in key cell-types like neurons.

In light of these challenges, large-scale assessment of gene regulation in brain tissues is necessary to better understand genetic and epigenetic features that govern brain biology and contribute to neurological disease. To achieve this goal, we have paired a well-phenotyped, high-quality brain tissue resource with production-scale, multi-omic profiling of multiple tissues from four brain donors. We generated occupancy maps of more than 100 TFs using chromatin immunoprecipitation followed by high-throughput sequencing (ChIP-seq). In addition to profiling bulk tissue from homogenates, we performed ChIP-seq in neuronal and glial cell types enriched with fluorescence activated nuclei sorting (FANS)^26, 27^. This approach improves the ability to associate epigenetic maps with their regulatory roles by reducing the confounding signal of multiple cell types mixed within homogenate tissues.

Here, we present a summary of our work with an emphasis on brain regions and cell types with the most thorough profiling and relevance to psychiatric disorders. By integrating the data produced here with published data, we describe the genomic localization of numerous TFs in relation to each other and their gene targets while identifying TFs whose occupancy patterns show increased heritability of several neuropsychiatric disorders.

## Results

### Experimental outline and genomic regions characterized by ChIP-seq

We performed genomic assays, including ChIP-seq, ATAC-seq, RNA-seq, phased whole genome sequencing, and DNA methylation profiling, on frozen postmortem brain tissue obtained from four donors that were free from and had no family history of psychiatric disorders (**Supplementary Table 1**). We performed assays on nine distinct brain regions; which included four large regions: dorsolateral prefrontal cortex (DLPFC), frontal pole (FP), occipital lobe (OL), and cerebellum (CB), as well as five smaller regions: anterior cingulate (AnCg), subgenual cingulate (SCg), dorsomedial prefrontal cortex (DMPFC), amygdala (Amy), and hippocampus (HC) (**Figure 1A**). We performed experiments with both homogenized tissue (“bulk”) and nuclei isolated using FANS (“sorted”), with 96 combined TFs and several histone marks (H3K4me1, H3K4me3, H3K27ac, H3K27me3, H3K9ac) assayed in large bulk samples and 76 TFs total in sorted nuclei from large regions (106 total TFs, detailed in **Supplementary Table 2**). By sorting with immunofluorescent labeling with NeuN and Olig2 antibodies, we were able to enrich for neurons (NeuN+), oligodendrocytes (Olig2+), and a mixed population of microglia and astrocytes (NeuN-/Olig2-) (**Figure S1A**). Representative ChIP-seq and ATAC-seq signals in sorted nuclei showed enrichment at promoters of marker genes of their respective cell type (**Figure S1B**). We prioritized TFs for ChIP-seq by selecting a large set for which there were validated ChIP-grade antibodies from the ENCODE Consortium^28^, many of which were done in our lab, as well as those with sufficient expression in the brain, and some level of established association with neurodevelopmental or neuropsychiatric traits (**Supplementary Table 3**).

**Figure 1.**
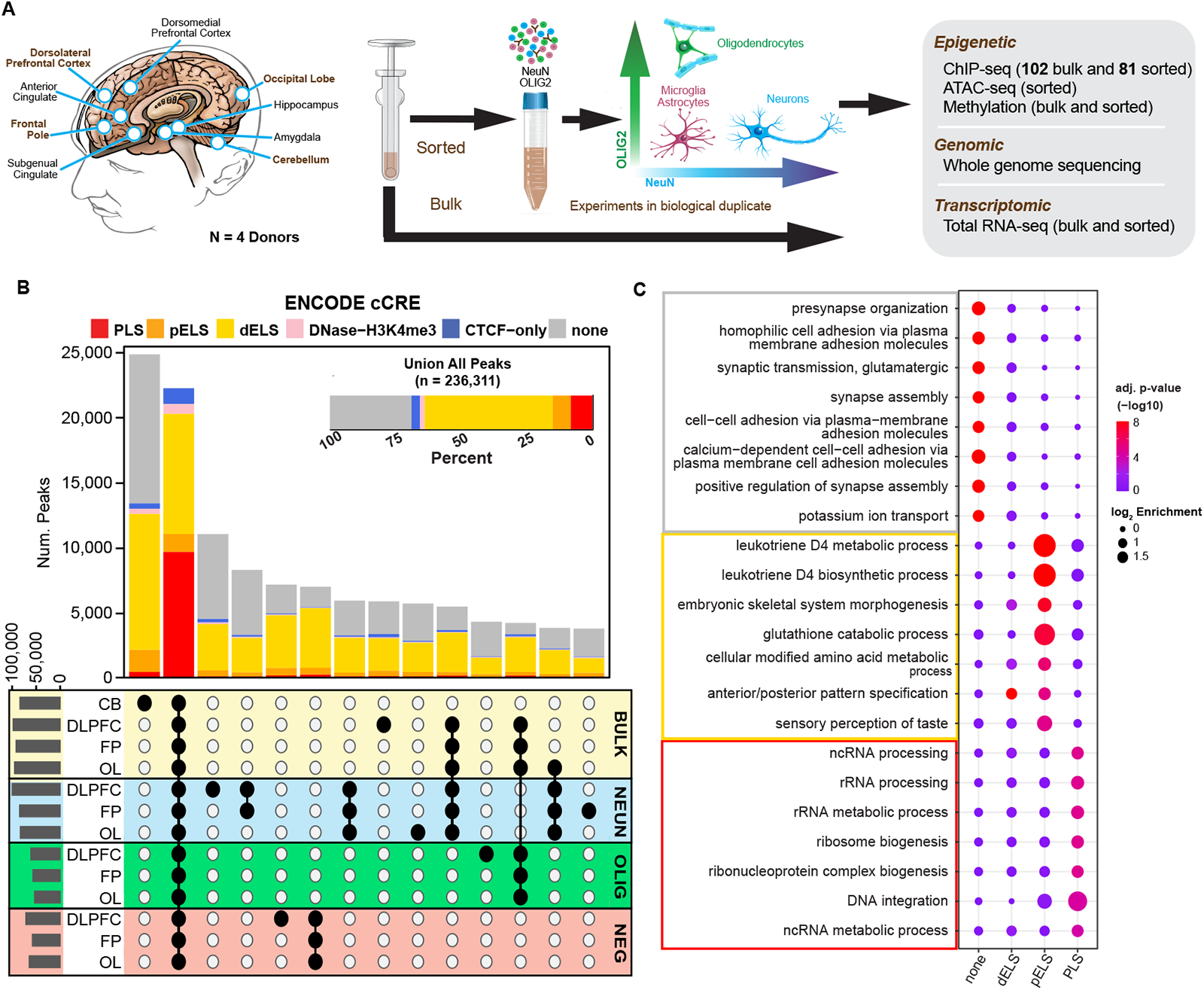
Overview of experimental design and profile of large brain regions tested. **A)** Left: Diagram of brain regions tested from four donors; large regions are in bold. Middle: Cell types used to perform experiments. Right: list of experiments. **B)** Intersections of peaks identified from TF ChIP-seq experiments from large regions (excluding histones). Barplot color indicates ENCODE cCREs annotation; “none” category indicates no annotation in cCRE registry. Upper barplot is the union of all TF peaks. **C)** GREAT enrichment of Gene Ontology terms for genes associated with peaks from the largest cCRE categories from the union of TF ChIP-seq experiments (top:none, middle: pELS and dELS, bottom: PLS).

Here, we primarily focus our analysis on the four large brain regions as they were the most extensively assayed. By generating the union of unique peaks for all TFs (excluding histone marks) across all four of these regions, we identified 239,361 distinct regions, which span 6.94% of the hg38 genome reference sequence, that are bound by at least one TF (**Figure 1B, Supplementary Table 4**). For each tissue type, 54% – 85% of regions are occupied by five or fewer TFs (**Figure S1C**). We categorized the regions of each intersection using the Registry of candidate *cis*-Regulatory Elements (cCREs) from the ENCODE Consortium^29^, and found that 49% were classified as distal enhancer-like sequences (dELSs) and 8% were classified as promoter-like sequences (PLSs, **Figure 1B**). Notably, promoters were shared across significantly more cell type/tissues than were dELSs (p*<*2.2e-16, Welch’s t-test), supporting the importance of enhancers in cell-type-specific functionality. The union of all identified regions revealed that 31% of all identified regions have no classification in the ENCODE Registry of cCREs, which to date consists of more than 1,500 cell lines and tissue types (Registry V3). Functional annotation of these regions with GREAT^30, 31^ revealed that the “none” category was enriched for neuronal pathways, while enhancers (“dELS” and “pELS”) and promoters (“PLS”) were enriched for more general gene sets (**Figure 1C**, **Supplementary Table 5**). We note that, while the majority of regions in the union are categorized as enhancers, there are several TFs whose peaks predominantly overlap promoters (**Figure S1D**).

### Correlation of ChIP-seq signal across TFs reveals co-association clusters

Many TFs have overlapping binding profiles and often cooperate to regulate gene expression^2, 32, 33^. We sought to identify correlations among TFs in our dataset to contextualize known TF interactions and discover new ones. To quantify TF associations within DLPFC-bulk experiments, we performed a principal component (PC) analysis on the signal (fold-change above background) of each TF at all regions identified by ChIP-seq. We then calculated the Pearson correlation between TFs using the first 20 PCs, accounting for 83.76% of the variation in ChIP-seq signals (**Figure 2A**, **Figure S2A and S2B**). PCs were used to quantify the relationship between TFs, as this approach allows us to assess subtle associations that might otherwise be masked by more dominant sources of variation, such as the unique binding pattern of cohesin-complex molecules (**Figure S2C**).

The resulting correlation heatmap can be segregated into three main clusters based on prominent features (**Figure 2B**). Cluster 1 appears to be composed of factors with relatively low preference for promoters, stronger average signal-to-noise ratios, and higher peak counts. In contrast, Cluster 3 is primarily composed of TFs that are rich in promoter binding, consistent with the observed overlap of H3K4me3 signal, and are generally anticorrelated with Cluster 1. Cluster 2 appears to be an intermediate cluster of factors that have some correlation across the full dataset. Similarities in binding among correlated TFs are evident when we examine factors spanning these clusters at a single locus (**Figure 2C**). As expected, we find the strongest signals at promoters (e.g. PKNOX1 and EGR1) and these are generally consistent among factors, even those from different clusters. In contrast, differences in binding are observed more frequently at distal sites, as we observed for CTCF and RAD21, members of the cohesin complex (**Figure 2C**). Similarly, members of the CTF/NF-I family NFIB and NFIC are correlated and share a unique peak at this example locus, although they are not found in the same cluster. Other notable correlations include EHMT2 (H3K9 methyltransferase, G9a) and REST (a neural gene repressor), which are known to participate together with chromatin remodeling complexes^34, 35^. Finally, we observe a correlation node within Cluster 2 involving TBR1 and SATB2, transcription factors with established roles in neuron development ^36, 37^. An axis of reciprocal correlations is formed between these two factors and 17 others, including SIN3B, ARNT, BCL11A, ZNF207, and ARID1B (**Figure S2D**). Although these factors are not widely known as direct interactors, these data suggest an association between these factors as they are all involved in neuron development^38–42^.

**Figure 2.**
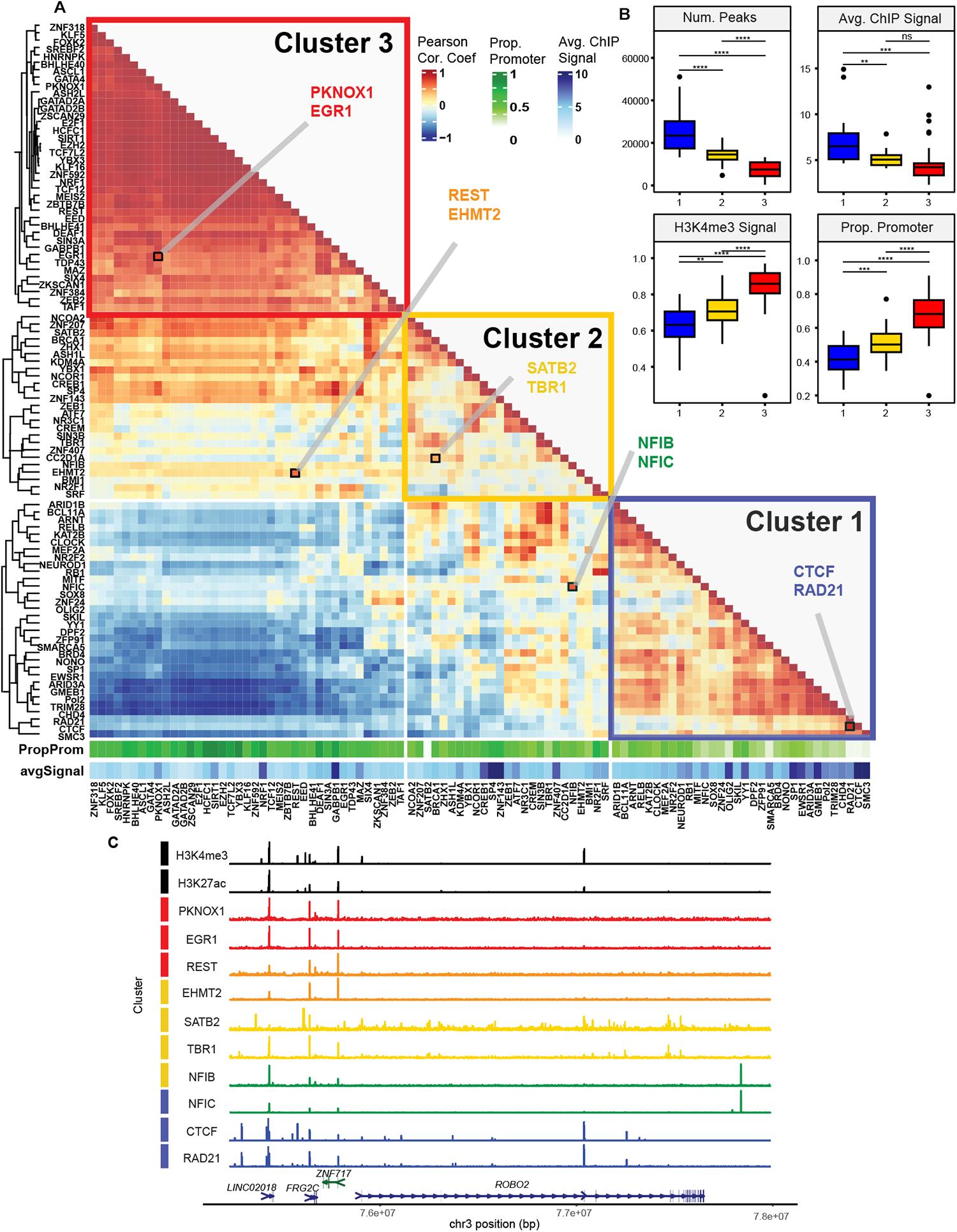
Comparison of ChIP-seq profile of TFs from DLPFC-bulk. **A)** Hierarchical clustering of TFs based on Pearson correlation of the first 20 PCs in DLPFC-bulk (k-means = 3). Noteworthy interactions within and between groups are highlighted. **B)** Quantification of key characteristics distinguishing the three main clusters (** indicates t-test p-value *<*0.01; *** p-value *<*0.001; **** p-value *<*0.001; n.s. non-significant). **C)** ChIP-seq signal tracks from highlighted TFs representing interactions between clusters along with histone marks H3K4me3 and H3K27ac. Gene track: small arrows denote directionality and large arrows denote possible transcript start sites.

### High-Occupancy Target sites can vary by factor preference and cell types

Although most regions identified across ChIP-seq experiments are bound by only a few factors (**Figure S1A**), many sites are occupied by a large number of TFs. Segments of hg38 with abundant binding of TFs are commonly referred to as High-Occupancy Target (HOT) sites and have been previously observed in various cell types^43–46^. These regions are highly permissive for TF binding and are enriched for non-specific binding of individual TFs. To define HOT sites within our data, we merged overlapping peaks from all experiments while setting a 2,000 bp threshold to prevent concatenation of adjacent regions. We ranked these regions by the number of unique TFs bound and set a threshold of the 90th percentile of the number of TFs to define HOT sites, similar to previous methods^43, 45^ (**Figure 3A**). When overlapped with ENCODE cCRE annotations, 66% of HOT sites are promoter-like sequences, particularly CpG island (CGI) promoters, while most non-HOT sites are distal enhancers (**Figure 3B**). To determine whether HOT sites are shared across tissues and cell types, we compared HOT sites identified separately in homogenate DLPFC, sorted neurons, oligodendrocytes, and astrocytes/microglia from DLPFC, and HOT sites defined in two cell lines from data available through ENCODE (**Figure 3C**). Cell type-specific HOT sites are largely distal enhancers, while those shared across cell types are mostly promoter-like sequences. HOT sites shared across all the cell types tested are heavily enriched for promoters of housekeeping genes (Odds Ratio = 8.245; p-value *<*2.2 x 10-16, Chi-squared test), which are known to coincide with HOT sites^43^.

We next looked at the binding profiles of individual TFs to measure their preference for co-localizing with other TFs and thus their tendency to be in HOT sites. This was done for each tissue/cell type by counting the number of TFs bound at each region in the union of all peaks, then measuring the proportion of peaks for increasing numbers of TFs bound. For example, SATB2 predominantly binds regions with few other TFs, while HCFC1 binds predominantly at HOT sites (**Figure 3D**). We measured the skewness of this distribution for each TF to quantify the propensity for binding at HOT sites (**Figure 3E**, **Supplementary Table 6**). Representative quantification in bulk DLPFC demonstrates the tendency of several factors to preferentially bind HOT sites and that the most HOT-skewed factors belong to the promoter-biased Cluster 3 in our PC analysis above (**Figure 2A**). Considering all tissue/cell type datasets, we found that skewness of HOT site binding is consistent between tissues (**Figure S3A**). Cortex-specific TFs SATB2 and TBR1 have negative-skewed profiles in the cortex regions, while NEUROD1, a neuronal differentiation factor, maintains this skewness across all three cortical regions and the cerebellum. Additionally, several TFs (e.g. PKNOX1) become more negatively skewed in neuronal nuclei relative to other tissue/cell types, while other TFs (e.g. NFIC) show the opposite trend (**Figure 3F and S3B**). We note that there is a positive relationship between the number of peaks called in an experiment and the calculated skewness, likely resulting from HOT sites and promoters being the majority of peaks identified in ChIP-seq experiments with lower signal-to-noise ratio (**Figure S3C**).

**Figure 3.**
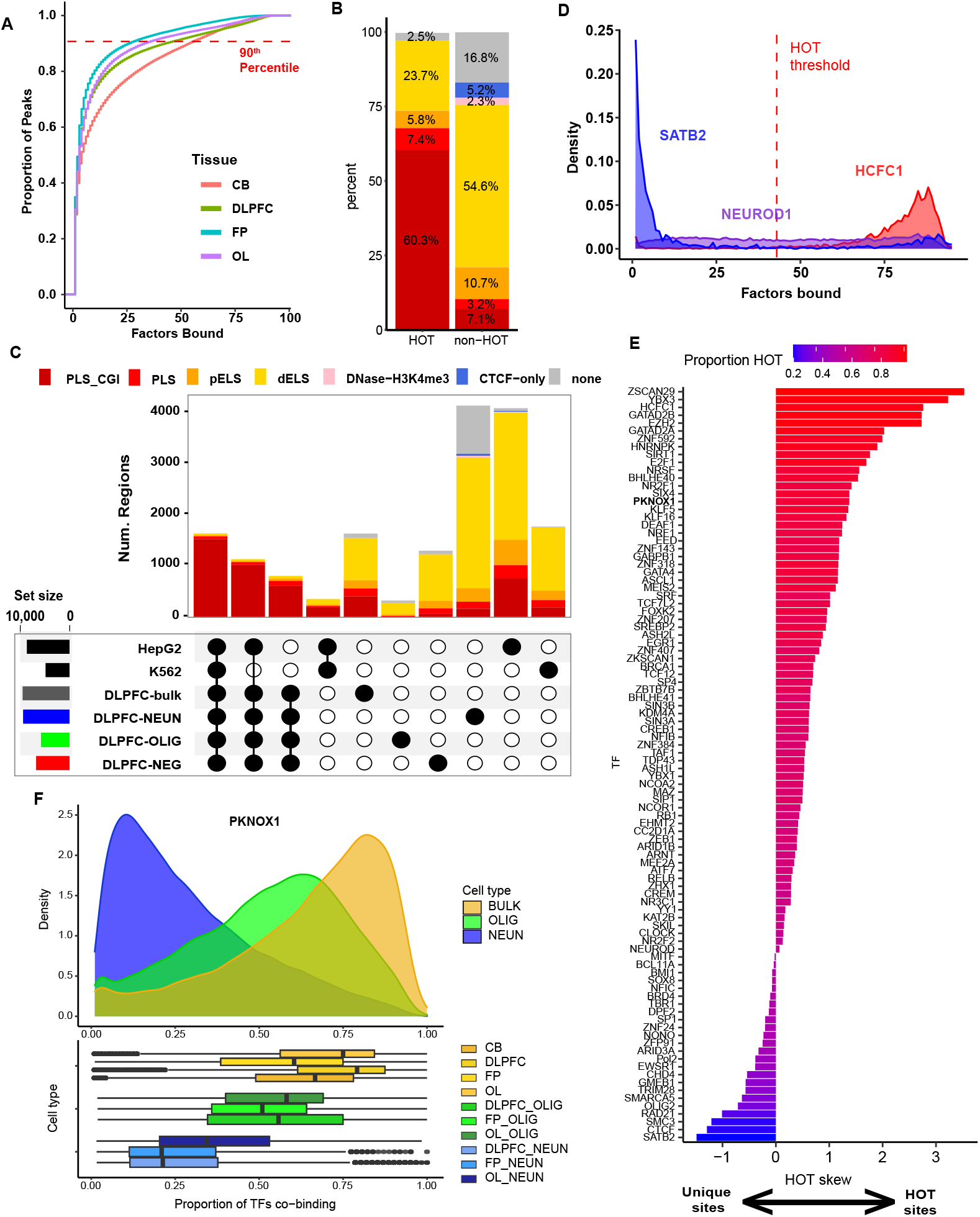
Identification and characterization of HOT sites. **A)** Cumulative distribution of peaks called from ChIP-seq of each large brain region demonstrating the cutoff for defining regions as “HOT”. **B)** Proportion of ENCODE cCREs in HOT and non-HOT DLPFC-bulk peaks. **C)** Intersections of HOT sites called from our ChIP-seq for all four cell-types of DLPFC and ENCODE ChIP-seq in cell lines HepG2 and K562. **D)** Distributions of peak occupancy for example TFs. **E)** Skewness values for the distribution of peak occupancy for all TFs in DLPFC-bulk. **F)** Distribution of peak co-occupancy in PKNOX1 across multiple tissues and cell types.

### Profiling enrichment and centrality of TF motifs

An important feature of many TFs is their ability to preferentially recognize and bind a particular DNA sequence motif. We compiled known motifs from the JASPAR 2022 database^47^ that correspond to TFs that we profiled, then measured the proportion of ChIP-seq peaks containing the motif and its centrality across all experiments in bulk DLPFC (**Figure 4A, Supplementary Table 7**,). Similar analyses of ENCODE data found a comparable percentage of peaks containing the expected TF motif (**Figure S4A**). 24 out of 49 factors had the highest relative proportion for their expected motif when scaled to all other factors in the dataset (**Figure S4B and C**). Some TFs are not the most enriched for their respective motif, but this is commonly due to closely related TFs having a stronger signal for the same motif (e.g., NFIB and NFIC, CREB1 and CREM). The centrality of all tested motifs, measured as the variance in distance from the motif to the center of the peak, showed a similar profile in that most TFs with high motif recognition also displayed high centrality for their expected motif relative to other TFs (**Figure S4D and E**). Plotting the normalized values together provides a visualization of the specificity of each factor for its expected motif in our experiments (**Figure 4A**). We also performed de novo motif discovery using MEME^48^. We found that TFs with higher specificity tend to match with their expected motif. We also note that peaks for a few TFs only revealed their expected motif once HOT sites were removed **(Figure S4B, Supplementary Table 8**). Previous ChIP-seq studies in human tissues^49^ have also found this to be true and it is likely driven by HOT sites composed of strong motifs for a few TFs^2^.

To identify commonly occurring motifs across experiments, we used a previously published approach that identifies and clusters motifs by similarity^50^. Among the 346 motifs that passed our quality metrics in experiments from bulk DLPFC, several motifs were repeatedly found among many experiments ((**Figure 4B**). These common motifs are highly GC-rich, emphasizing the role of GC-preferring factors, such as the NRF and SP families (represented in clusters 4 and 10 respectively), in regulating gene expression in the brain^51–53^. The largest motif cluster (1) resembles that of THAP11, which was previously shown to be common across ChIP-seq experiments and may be involved with promoter-promoter interactions^54, 55^. These motifs highlighted in Figure 4B are often enriched in de novo motif discovery across individual experiments.

Because HOT sites are regions that lack motif diversity, we calculated the difference in motif recognition between the full and HOT-depleted peak set for each TF and compared it to the GC content of the expected motif (**Figure 4C**). TFs for which the expected motif was more enriched in HOT sites tended to be those with higher GC content in the motif sequence, reflecting the common motifs highlighted in Figure 4B. Conversely, removing HOT sites improved enrichment for motifs with lower GC content. For example, when de novo motif calling was performed on the top 500 NFIB ChIP-seq peaks, the expected NFIB motif was the second ranked motif called by MEME, but after excluding HOT sites, it became the top ranked motif with drastically increased enrichment (**Figure 4D**) and sharpened centrality (**Figure S5A**). Compared with ATAC-seq peaks, ChIP-seq experiments from all tissues and cell types generally showed a greater enrichment for the expected motif (**Figure S5B**). Altogether, these data show that the ChIP-seq experiments performed in this study show enrichment for their respective motifs by multiple metrics and that the results are influenced by the GC content, likely due to its association with HOT sites^43^.

**Figure 4.**
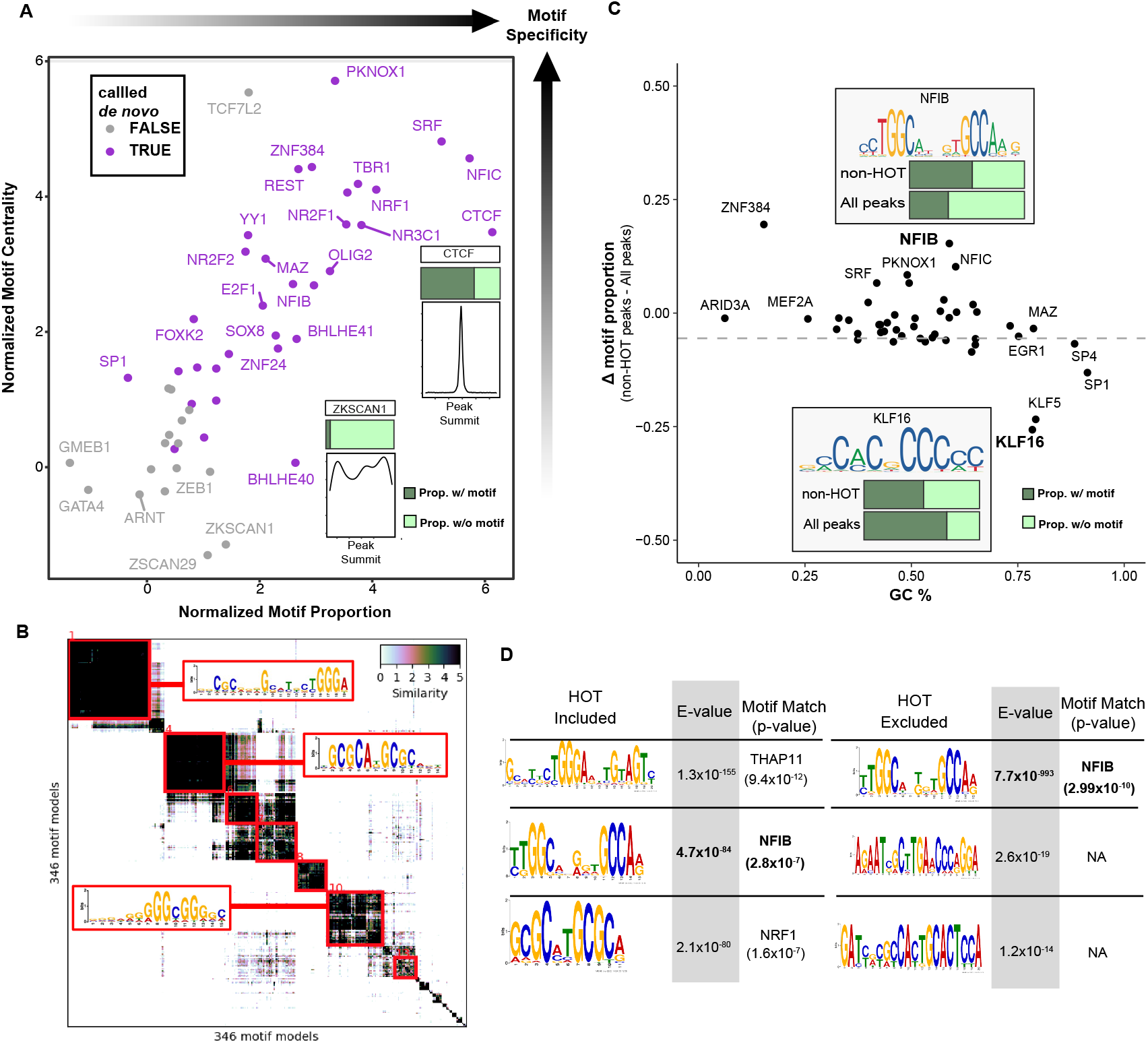
Motif recognition by transcription factors. **A)** Schematic demonstrating the specificity of individual TFs for their expected motif relative to the other TFs tested which had a matching motif in the JASPAR 2022 database (49 in DLPFC-bulk ChIP-seq). Motif Proportion is the scaled proportion of peaks for individual TFs containing their expected motif relative to other TFs tested. Similarly, Motif Centrality is the scaled centrality of the expected motif within respective peaks relative to other TFs tested. Increase in these two metrics indicate a ChIP-seq that is more specific for its expected motif relative to other ChIP-seq experiments. **B)** Matrix of de novo motif calls using top 500 peaks from all ChIP-seq experiments in DLPFC-bulk. The most prominent clusters are highlighted along with the called motif. **C)** Dot plot showing the relationship between motif GC content and the difference in proportion of peaks with expected motif matches found in all peaks from a ChIP-seq experiment and the proportion with HOT sites removed. **D)** NFIB ChIP-seq as an example of the effect that removing HOT sites has on de novo motif calling using MEME-ChIP. The left table uses the top 500 peaks ranked by q-value; the right table uses the top 500 peaks excluding HOT sites. E-value enrichment for each called motif is calculated by MEME and highlighted in gray. TOMTOM calculated the p-value of the match between the de novo motif and the corresponding position weight matrix from the JASPAR database.

### Ability of predictive models to predict ChIP-seq results

With the growing availability of TF-occupancy data from sources such as the ENCODE Consortium, there have been efforts to predict TF binding with computational models trained on existing data. To test predictions versus our experimental results, we compared our ChIP-seq datasets to models generated by Virtual ChIP-seq^56^, a peak predictor that is trained on many datasets from CCLE ^57^ and ENCODE ^28^. There were 16 TFs in both our dataset and the Virtual ChIP-seq trained models that used multiple cell lines for each TF. We assessed the precision-recall at increasing posterior probability cutoffs calculated by Virtual ChIP-seq and derived the area under the precision-recall curve (auPR) for each TF (**Figure 5A, Supplementary Table 9**). Where applicable, we also assessed the precision-recall of ATAC-seq peaks containing the expected motif to gauge its predictive accuracy alongside the respective model (**Figure 5A**, red asterisk). Higher auPR, as demonstrated by CTCF and NRF1, is indicative of stronger agreement between the predictive model and the experimental data. Models with relatively low auPR, for factors such as EGR1 and REST, failed to outperform ATAC-seq peaks containing motifs for a given level of recall. We also calculated the auPR on ENCODE datasets for the GM12878 cell line (included in the training data) and found that the auPR for predictions using data from bulk DLPFC were generally comparable (r = 0.69, p-value = 0.0093) to those of the cell lines (**Figure 5B**). We replicated this approach in sorted nuclei and found the results to be consistent across multiple cell types, with predictive models typically having higher auPR for experiments in bulk tissues compared to NeuN+ and Olig2+ sorted nuclei (**Figure S6B**). However, we note that the predictive power of these models is primarily derived from HOT sites, as the auPR is drastically reduced across most models when HOT sites are removed (**Figure 5C**). This is likely due to the preferential removal of promoter regions, which are inherently correlated with expressed genes and open chromatin. As CTCF is not typically associated with promoter regions, it is the least affected by HOT site removal. We calculated the enrichment for cCRE categories of predicted peaks that matched the experimental data and show that distal elements are depleted, further exemplifying the bias of a predictive model for predicting peaks in promoters (**Figure 5D**).

**Figure 5.**
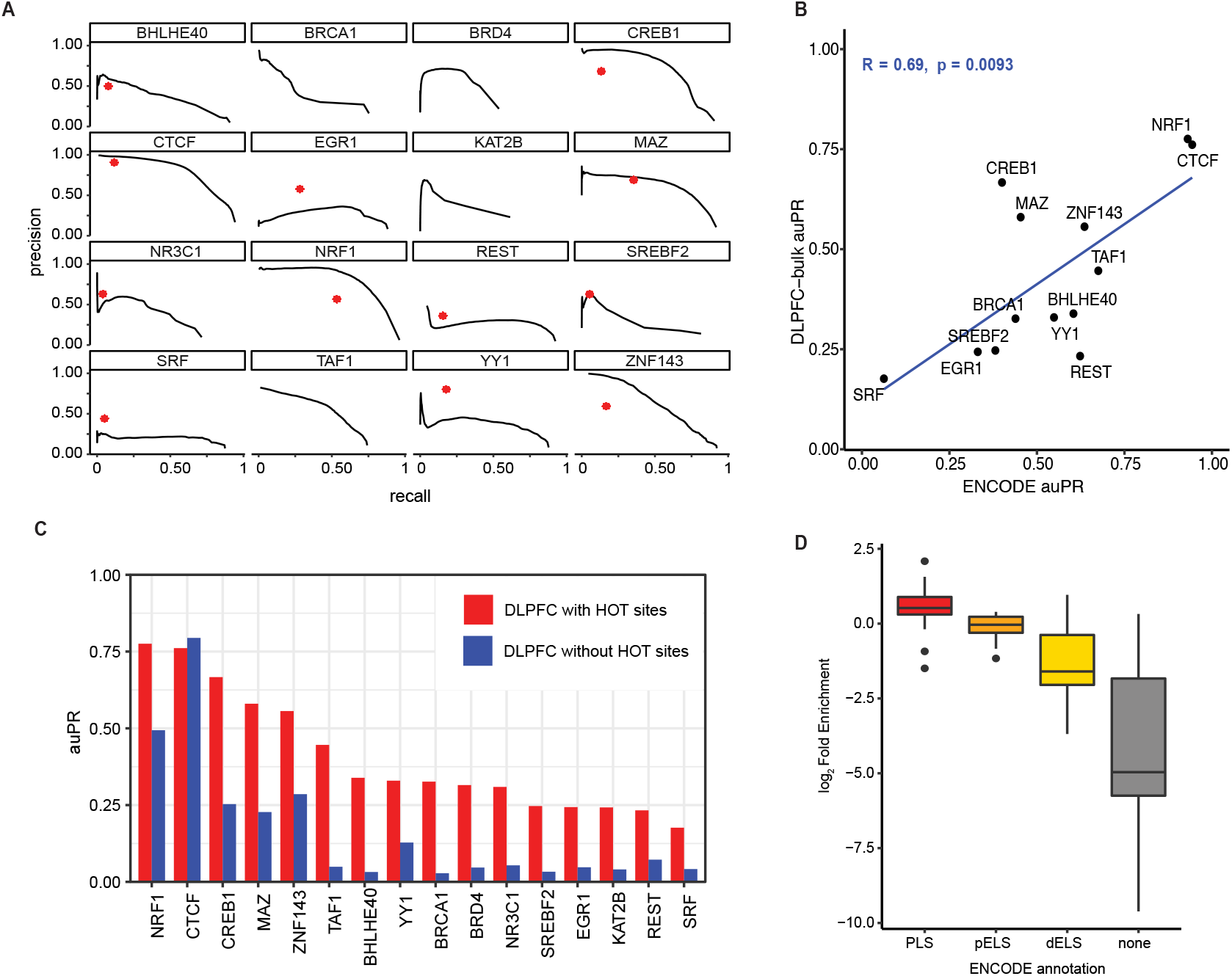
Comparison of ChIP-seq results with predictive models. **A)** Precision-recall curves of predictions made by Virtual ChIP-seq compared to ChIP-seq results. Red asterisk represents precision-recall of ATAC-seq peaks containing the expected motif. **B)** Calculated Virtual ChIP-seq predictions made using ChIP-seq from our experiments using DLPFC-bulk or from experiments on GM12878 available through ENCODE database (R = Pearson correlation coefficient). **C)** auPR with and without HOT sites. **D)** The enrichment for cCREs categories in regions predicted at a posterior probability *>*0.5 over unpredicted regions.

### Multi-omic Integration

We classified genes as to whether or not their promoters overlap a HOT site. Genes with HOT-site promoters have higher median expression and stronger indicators for active gene expression, including H3K4me3, chromatin accessibility, and hypomethylation (**Figure 6A**). We quantified the functional consequence of TF localization on gene regulation by correlating their binding with gene expression using data from NeuN+ sorted DLPFC nuclei. To measure the impact of individual TFs on gene expression, we implemented a linear model for each factor to estimate the effect on gene expression while controlling for the number of TFs bound at the promoter (**Figure 6B**, left). Most TFs have a positive impact on gene expression with the largest effect sizes measured in genes with non-HOT promoters, suggesting that individual TFs have a relatively small impact at HOT-promoters. Genes with HOT promoters appear to have become saturated, as there is no association between activation signals such as ATAC-seq or TF count and transcription (Figures S7A & B). We performed a similar analysis using ChIP-seq signal at distal elements that were linked to specific genes in our previous single-cell study using 10x Genomics multiomic technology^58^ (**Figure 6B**, right), and demonstrated a similar trend, albeit with smaller effect sizes overall. These findings were replicated in bulk DLPFC samples, although there are discrepancies in some TFs having a strongly negative effect on expression likely caused by differences in gene expression between cell types (**Figure S7C**).

We determined the cell type composition of TF peaks from bulk DLPFC ChIP-seq by overlapping with ATAC-seq peaks from sorted nuclei. We identified TFs that were biased towards specific cell types, notably SATB2, TBR1, and BCL11A for NeuN+ nuclei, OLIG2 and SOX8 in Olig2+ nuclei, and MITF for NeuN-/Olig2-(**Figure 6C**). These results were replicated by measuring the enrichment of these factors for cell type-specific linkages from our single-nuclei multiomic data (**Figure S7D**), of which all were shown to significantly affect gene expression in the linear models for either sorted or bulk ChIP-seq. We note that TFs in bulk OL have a greater proportion of peaks exclusive to NeuN+ ATAC-seq (**Figure S7E**), likely due to the higher proportion of neurons in OL relative to DLPFC as observed from FANS (**Figure S1A**). Of all TFs tested in bulk DLPFC, ATAC-seq signal in sorted nuclei was disproportionately high across regions bound by each of the three neuronal TFs when corrected for the relative number of total TFs bound (**Figure 6D**). This effect is not observed in the corresponding sorted ATAC-seq signal in other cell type-enriched factors such as OLIG2 and MITF, suggesting that this effect is neuron-specific.

**Figure 6.**
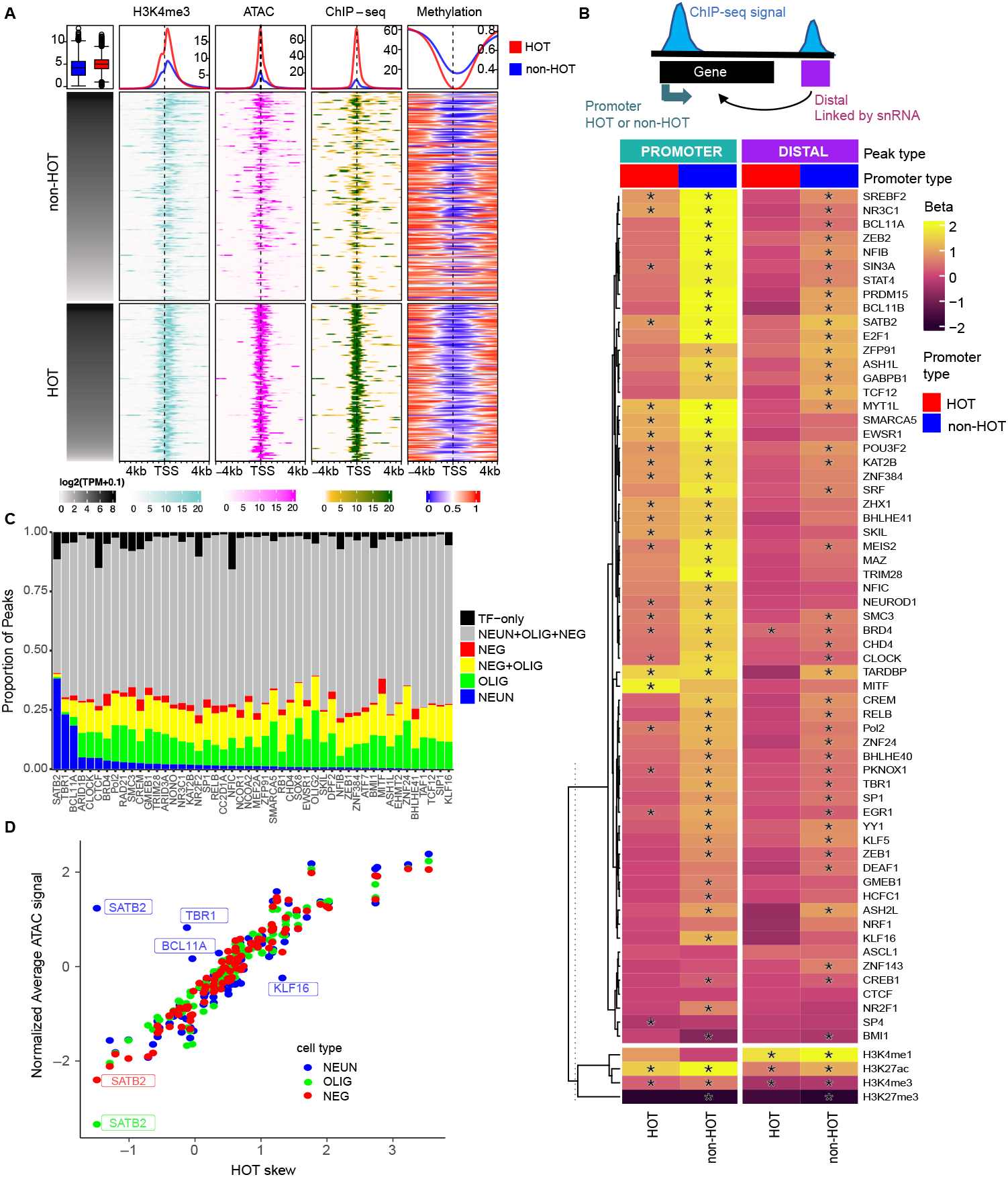
Correlating TF binding with gene expression and chromatin accessibility. **A)** Heatmap demonstrating the relationship between gene expression and markers for active gene expression and gene promoters such as H3K4me3, chromatin accessibility (ATAC-seq), cumulative TF binding (ChIP-seq), and methylation in experiments from DLPFC-bulk. Cumulative ChIP-seq signal was generated by combining peaks called from all TF ChIP-seq experiments in DLPFC-bulk into a single bed file and converting it into a bigwig file. Methylation signal was derived from whole genome bisulfite sequencing. **B)** Calculated beta values from linear models estimating the effect that each TF has on predicting gene expression based on ChIP-seq signal in DLPFC-NEUN at the promoter (left) or a linked distal region (right) while controlling for the number of co-bound TFs. Top: Illustration of the ChIP-seq signal at promoters and distal sites linked to gene expression. Bottom: beta values for individual TFs with histone marks and ATAC-seq shown separate. **C)** ChIP-seq peaks of TFs in DLPFC-bulk segmented by their overlap with ATAC-seq from sorted nuclei. Shown here are TFs with the highest proportion of non-Common peaks. **D)** Plot of average ATAC-seq signal (normalized across cell-types) across ChIP-seq peaks for each TF from DLPFC-bulk. HOT skew is calculated as in Figure 3. NEUN: NeuN+ nuclei; OLIG: Olig2+ nuclei, NEG: NeuN-/Olig2-nuclei.

### Enrichment of heritability for psychiatric disorders within TF peaks

To identify disease-relevant TFs, we integrated our genomic data with GWAS findings using stratified LD score regression (sLDSC), a statistical method developed to estimate and partition SNP heritability of a trait by different functional genomic annotations^59^. We used the ChIP-seq and ATAC-seq datasets performed in bulk tissue and sorted nuclei along with GWAS summary statistics from various neuropsychiatric and neurodegenerative diseases.

Using sLDSC analysis for experiments from DLPFC, there is a clear increase in the enrichment of neuropsychiatric disease markers for experiments from NeuN+ sorted nuclei compared to those from bulk and other sorted cell types (**Figure 7A**, Supplementary Table 10). This is evidenced by the increase in the number of individual TFs significantly associated with neuropsychiatric disorders and overall increase in enrichment. While histone marks and ATAC-seq regions have similar enrichment between bulk and NeuN+ datasets, the individual TFs show a clear increase in enrichment for associated diseases. By looking at a single factor (PKNOX1), we highlight the increase in enrichment provided by ChIP-seq compared to ATAC-seq (**Figure 7B**). Neuronal TFs TBR1 and BCL11A were significantly enriched for schizophrenia and bipolar disorder in both bulk and NeuN+, but SATB2 is not. We note an enrichment of Alzheimer’s disease risk alleles for several TFs in the NeuN-/Olig2-nuclei, notably for REST in the DLPFC and FP (**Figure 7A, S8A**). TFs were generally not enriched in experiments from Olig2+ nuclei. No dataset from this study was significantly enriched for non-neuronal phenotypes that served as negative controls, including immune diseases or other physical traits. However, TFs from this study that were also profiled by ENCODE in the GM12878 immune cell line did show enrichment in those experiments for inflammatory disorders, highlighting the cell type specificity of DNA binding by these factors (**Figure S8B**).As an example, we highlight a genomic region where a PKNOX1 peak from NeuN+ DLPFC nuclei overlaps a risk locus for schizophrenia^14^ (**Figure 7C**). The most significant SNPs were in the intronic regions at the 3’ end of *TSNARE1* (bottom). Overlapping PKNOX1 peaks with linkages from our previous single-cell multiomics study shows that this particular region is correlated with *ADGRB1* and *ARC* expression in neurons, two genes that were previously implicated in schizophrenia^60–62^. We note that this link does not overlap an ATAC-seq peak, although it was identified as an ATAC-seq peak in the single-cell study and shows some signal above background in our study.

**Figure 7.**
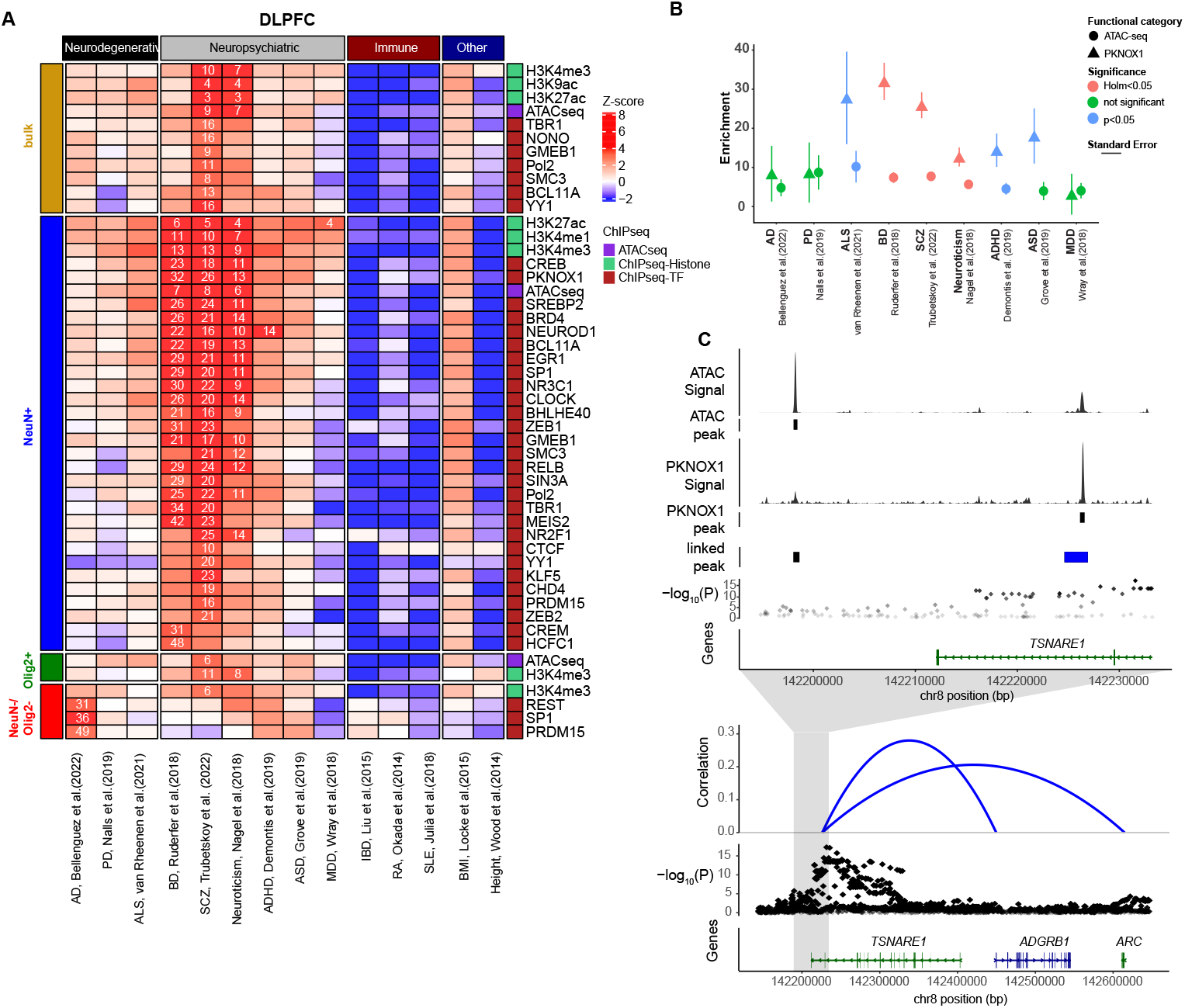
Association of TFs with disease through GWAS traits. **A)** Results from sLDSC measuring heritability of TFs with GWAS traits grouped by disorder type and separated by cell type. Heatmap indicates coefficient Z score from sLDSC of TF ChIP-seq peaks from each cell-type combined with 97 baseline features. Feature-trait combinations with a Z score significantly larger than 0 (one-sided Z test with alpha = 0.05, p values corrected within each trait using Benjamini-Hochberg method) are indicated with a numeric value reporting the enrichment score. **B)** Direct comparison of sLDSC results for single TF (PKNOX1) with ATAC-seq to highlight the increase in heritability enrichment for neuropsychiatric disorders. **C)** Browser tracks of a region containing SNPs with statistically high association with schizophrenia (Trubetskoy et al 2022). Top: Zoomed in region to show signal and called peaks from both ATAC-seq and PKNOX1 ChIP-seq in relation to significant peaks from GWAS and links, which are regions identified by single-cell multiomic study in cortex tissue identifying differentially accessible regions significantly associated with nearby genes. Bottom: Zoom out of region. Arcs represent the correlation between the linked-peak overlapping a PKNOX1 peak in the top panel with nearby genes *ADGRB1* and *ARC*.

## Discussion

Large epigenetic datasets, such as those provided by ENCODE, have been cited in thousands of publications and have played a major role in understanding the role of TFs in gene regulation^25^. However, such datasets typically focus on cell lines that have limitations with respect to modeling gene regulation observed in human tissues, particularly brain. Recent studies have highlighted the importance of using human tissues for studying epigenetic regulation in normal brain function and understanding of psychiatric disorders^17, 18, 23, 24^, but such studies often focus on general markers such as histone modifications and chromatin accessibility. Here, we have used ChIP-seq to map the binding of more than 100 proteins that regulate chromatin structure, including histones, transcription factors, and chromatin remodelers (**Figure 1**). ENCODE has generated an extensive registry of 1,063,878 candidate *cis*-Regulatory Elements (cCREs) defined from more than 1,500 cell lines and human tissues, of which 235 are considered neural (defined as brain, spinal cord, or nerve)^29^. Nearly 30% of the regions identified by TFs here are not annotated in this cCRE registry and are enriched for neuronal pathways, highlighting the unique attributes of our study.

A benefit to performing ChIP-seq experiments on such a large number of TFs in uniform contexts is that it allows the study of TFs as a group, including the identification of HOT sites. We identified many HOT sites outside of typical CpG islands, gene promoters, and sites found in ENCODE data (**Figure 4C**), emphasizing the need to experimentally determine such sites in relevant tissues and cells. Providing this information will aid researchers in using this resource and potentially help to interpret analyses from other studies by identifying those TFs and regions that exhibit unique occupancy patterns. For example, knowing which TFs are most selective for specific sites (e.g., **Figure 3C**) may help in prioritizing factors in future studies. Similarly, knowing regions of seemingly indiscriminate binding can identify genes whose expression is relatively unchanged regardless of which TFs are bound at the promoter or potential enhancers (**Figure 5**). Analyzing data from regions outside of HOT sites also substantially improves de novo motif discovery for some factors (**Figure 4C**).

As the generation of epigenetic and multi-omic datasets of brain tissue increases, there will be a greater need for resources capable of predicting TF-binding at sites of interest. The ability to overlap findings with ChIP-seq from human samples will be a valuable resource that is difficult to replicate in silico. For example, comparison of predicted TF-binding results from Virtual ChIP-seq to experiments from our study showed major discrepancies that suggest limitations to the accuracy of computational models that rely solely on motifs and ATAC-seq for predicting the function of many TFs (**Figure 5**). Most of the predictive power from these models come from predicting HOT sites (which are more easily predicted and the most well-understood CREs) because the auPR plummets upon their removal. We showed that the “none” cCRE category lacking annotation in the ENCODE registry is enriched for neuronal pathways (**Figure**1**C)** and is the most depleted cCRE category in the predictions (**Figure 5D**). These findings accentuate the unique value of the brain-derived data described here which may improve model performance in the future.

Analyzing ChIP-seq data with sLDSC connects specific TFs to their potential roles in psychiatric disorders using GWAS data from neurological diseases. The increased enrichment of heritability for TFs in specific cell types (psychiatric in neuron-enriched and Alzheimer’s in microglia-enriched) relative to histone marks and ATAC-seq shows the value of performing ChIP-seq in disease-relevant cell types (**Figure 7**). Several highly enriched TFs have been previously implicated in psychiatric disorders, such as PKNOX1^63^,CREB1^42^,EGR1^10^, and NEUROD1^65^. Using PKNOX1 as an example, we show binding at a region of significance for schizophrenia that would likely be missed in ATAC-seq analysis and is associated with neuronal expression of multiple disease-related genes. This dataset provides a multitude of opportunities for functional annotation and analysis of disease-associated genes and CREs.

To date, this is the largest study directly measuring TF binding in human postmortem brain and is complementary to other large-scale genomic efforts (e.g., ENCODE, PsychENCODE, GTEx, etc.). These data and the accompanying analyses will serve as a resource to understand genome regulation in psychiatric diseases and will be publicly available through the PsychENCODE Consortium and available for download at the following link: https://doi.org/10.7303/syn4921369.

## Acknowledgments

This study was supported by NIH grant 5R01MH110472 awarded to R.M.M. and G.M.C, the Memory and Mobility Fund from HudsonAlpha Institute for Biotechnology, and support from the Pritzker Neuropsychiatric Research Consortium. We especially thank the brain donors and their families without whom this research would not have been possible.

## Author contributions

Conceptualization: R.M.M, G.M.C., J.M.L. Investigation: J.M.L., L.R., I.R.N., K.T.L., R.J., Data Curation: J.M.L., L.R., A.G.A. Formal analysis: J.M.L., A.G.A., B.M., I.R.N. Provision of brain tissue and resources: W.E.B, B.G.B, P.C., A.S., S.J.W., H.A. Writing – Original Draft: J.M.L. Funding acquisition: R.M.M., G.M.C.

## Data and code availability

The data will be publicly available through the PsychENCODE Consortium and available for download at the following link: https://doi.org/10.7303/syn4921369. All supplementary tables are available by request. All original code generated during this study is publicly available at github.com/aanderson54/Loupe_BrainTF and has been depositied at Zenodo (DOI: 10.5281/zenodo.8065798).

## Methods

### Brain tissues

Human brain tissues were obtained from collaborators at the Department of Psychiatry and Human Behavior, University of California Irvine; dissections were performed at the Molecular & Behavioral Sciences Institute, University of Michigan. Both donors died without prolonged agonal state and had no personal or family history of psychiatric disease. Samples were stored at -80*^◦^*C.

### Chromatin Preparation: Bulk

Buffers required: RIPA: 1x PBS (Cytiva SH30256.02), 1% NP-40, 0.5% sodium deoxycholate, 0.1% SDS. Farnham Lysis buffer: 5 mM PIPES pH 8.0 / 85 mM KCl / 0.5% NP-40. Roche Protease Inhibitor Cocktail Tablet (Sigma 11836145001 for 50 ml or mini tablets 11836153001 for 10 ml).

In order to reduce variability between experiments and reduce sample loss of precious tissue, chromatin inputs for ChIP-seq were prepared in large batches (enough for 20-100 experiments for small and large regions respectively). Techniques for generating chromatin from frozen human tissue were similar to those described previously from our lab64. For processing frozen brain tissue, all instruments and materials are chilled on dry ice and kept cold until crosslinking. To begin, a small portion of each dissected brain region is broken off using a metallic block and hammer with care taken to obtain an even balance of gray and white matter when possible. The starting material for each region (2500mg for large regions and 500mg for small regions) was sealed inside of a Covaris Tissue TUBE (Covaris 520021, 520023) and thoroughly pulverized using the chilled hammer and metallic block, with intermittent submerging into liquid nitrogen to maintain cold temperature and brittleness. At this point, a small amount of pulverized tissue (*<*100mg) is preserved in an Eppendorf tube at -80*^◦^*C to be used in parallel experiments. The remaining tissue can be used immediately or stored for a few days at -80*^◦^*C.

For crosslinking, pulverized tissue was poured into a conical tube holding PBS plus protease inhibitor at room temperature and a small volume of PBS is used to wash the remaining contents of the Covaris bag. Immediately, add 37% formaldehyde (Sigma F8775-25ML) to a final of 1% formaldehyde and rotate end-over-end for 10 minutes at room temperature. Fixation was halted by adding 2.5M Glycine to a final concentration of 0.125M and rotating for another 5 minutes at room temperature. Pour the fixed tissue mixture into a Dounce homogenizer and pass the loose pestle 5 times to break up any remaining large pieces. Spin the mixture in a conical tube at 5,000g for 5 minutes at 4*^◦^*C. The supernatant is discarded and the pellet is resuspended and washed in cold PBS. Spin and 5,000g for 5 minutes at 4*^◦^*C. Wash twice more with cold PBS. Resuspend final pellet in Farnham Lysis buffer plus protease inhibitor and hold on ice for 5 minutes. Pour contents in Dounce homogenizer on ice and complete 10 pulses with tight pestle. Pour into new conical tube and spin 5,000g for 5 minutes at 4*^◦^*C. Resuspend pellet in cold RIPA buffer plus protease inhibitor. Sonicate tissue using Diagenode Bioruptor Pico (10 cycles; 30 seconds on/ 30 seconds off) using 1.5mL microtubes (Diagenode C30010016) with 300uL per tube (several rounds needed). Pool the sonicated chromatin into a conical tube and centrifuge 16,000g for 30 minutes at 4*^◦^*C to remove any insoluble debris. Collect supernatant, raise to final volume with RIPA, and dispense working aliquots into 1.5mL Eppendorf tubes held on dry ice. Store at -80*^◦^*C.

### Chromatin Preparation: Sorted

Buffers required: Nuclei Extraction Buffer (NEB): 0.32 M Sucrose, 5 mM CaCl2, 3 mM Mg(Ac)2, 0.1 mM EDTA, 10 mM Tris-HCl, 0.1 mM PMSF, 0.1% Triton X-100, 1 mM DTT. Before use, add protease inhibitor cocktail according to manufacturer recommendation. Sucrose Cushion Buffer (SCB): 1.6 M Sucrose, 3 mM Mg(Ac)2, 10 mM Tris-HCl, 1 mM DTT. Interphase Buffer: 0.8 M Sucrose, 3 mM Mg(Ac)2, 10 mM Tris-HCl. Blocking buffer: 1x PBS, 1% BSA (Sigma 3117332001), 1 mM EDTA. Pellet buffer: add 200 uL 1 M CaCl2 to 10 mL SCB.

Methods for extracting and sorting nuclei from postmortem brain are similar to previously published methods67. Here, approximately 500 mg of tissue was placed into a chilled 40 mL Dounce homogenizer containing 5 mL of NEB on ice and allowed to partially thaw to ease douncing (2-3 minutes). Extract nuclei by douncing with “tight” pestle 30-40 times until the tissue is homogenized. Transfer to a 15 mL conical tube on ice, wash glassware with 5 mL NEB and add to 15 mL tube. Fix chromatin by adding 625 uL of 16% formaldehyde (methanol free, Thermo 28906) to a final concentration of 1% and rotate end-over-end at room temperature for 10 minutes. Halt fixation by adding 500 uL of 2.5 M Glycine and incubate another 5 minutes rotating at room temperature then place homogenate back on ice. During fixation, prepare sucrose gradient in 2 ultracentrifuge buckets (Beckman Coulter cat: 344058) by layering 5 mL of Interphase buffer atop 10 mL of SCB in each. Carefully layer nuclei homogenate atop sucrose gradient, balance with NEB, then ultracentrifuge at 24,000 rpm for 2 h using SW28 swinging bucket rotor (Beckman Coulter). Upon completion, inspect tubes for a visible pellet of nuclei at the bottom of the tube. Remove debris at interphase first by using a 25 mL graduated pipette, then continue removing the remaining sucrose gradient being careful not to disturb the nuclei pellet. Carefully resuspend the pellet in 1 mL cold PBS and transfer to a 15 mL lo-bind tube containing 2 mL PBS on ice. Wash ultracentrifuge tubes with 1 mL cold PBS and combine in 15 mL tube to a final volume of 10 mL, inverting to mix. Centrifuge the nuclei at 1,000xg for 10 minutes at 4*^◦^*C to remove residual sucrose. Label nuclei by resuspending pellet in 5 mL blocking buffer with NeuN-488 antibody (Millipore MAB377X) and Olig2 antibody (Abcam ab109186) at 1:5,000 each. Incubate nuclei in staining buffer with rotation for at least 1 hour at 4*^◦^*C. Spin nuclei 500 x g for 5 minutes to pellet, remove supernatant, then resuspend in 5 mL blocking buffer with goat-anti-rabbit-647 (ThermoFisher A-21245) at 1:5,000 and DAPI at 1:100,000. Incubate for at least 1 hour at 4*^◦^*C with rotation (overnight preferred). Remove stain by centrifuging 500xg 5 minutes at 4*^◦^*C and resuspending in 3 mL cold PBS. Hold on ice and proceed immediately to sorting.

Nuclei were sorted using Sony MA900 with a 70 um nozzle and pressure not exceeding pressure setting of 7. Gates were set to capture those populations that were positive for 488 signal (NeuN+), positive for 647 signal (Olig2+), or negative for both (NeuN-;OLIG-). Each population was collected into 5 mL tubes held at 4*^◦^*C and pooled into 15 mL lo-bind tubes on ice. Purity of selected samples were typically ¿95% based on reanalysis of sorted samples. To concentrate nuclei for downstream analysis, add approximately 2 mL of pellet buffer per 10 mL of sorted nuclei and rotate at 4*^◦^*C for 15 minutes. Centrifuge 500 x g for 10 minutes at 4*^◦^*C, after which a pellet should be visible. Remove supernatant and carefully resuspend pelleted nuclei in at least 3 mL cold PBS to wash. Centrifuge 500xg for 5 minutes at 4*^◦^*C.

To generate chromatin for ChIP-seq, resuspend pellet in cold RIPA plus protease inhibitor at approximately 3 million nuclei per 250 uL. Transfer 250 uL of each sample to the Bioruptor tubes and sonicate tissue using a Bioruptor Pico (8 cycles; 30 seconds on/ 30 seconds off). Pool the sonicated chromatin into a 1.5 mL DNA lo-bind conical tube and centrifuge 12,000xg for 5 minutes at 4*^◦^*C to remove any insoluble debris. Collect supernatant into a separate tube, add RIPA to final volume equivalent to 500,000 nuclei per 100 uL, then dispense working aliquots into 1.5 mL tubes held on dry ice. Store at -80*^◦^*C.

### ChIP-seq

Buffers required: BSA wash: 1x PBS + 5 mg/mL BSA. LiCl wash: 100 mM Tris at pH 7.5, 500 mM LiCl, 1% NP-40, 1% sodium deoxycholate. TE buffer: 10 mM Tris-HCl at pH 7.5, 0.1 mM Na2EDTA. IP elution buffer: 1% SDS, 0.1 M NaHCO3.

Antibodies used in ChIP-seq are detailed in Supplementary Table 3.

Protocols for ChIP-seq are similar to those for frozen tissue previously described by our lab^66, 68^ and consistent with techniques recommended by the ENCODE consortium (www.encodeproject.org/documents). Briefly, Dynabeads (ThermoFisher Scientific; 11201D anti-mouse or 11203D anti-rabbit) were washed with cold 1x PBS + 5 mg/mL BSA then combined with respective antibodies at a desired ratio of 5 ug antibody to 150 uL Dynabeads (dependent on concentration from manufacturer) at a final volume of 200 uL and held at 4*^◦^*C overnight with rotation. The following morning, tubes of aliquoted chromatin are thawed on ice and bead/antibody complex is washed with PBS + 5 mg/mL BSA solution. Beads are ultimately resuspended in 100 uL RIPA and brought to 200 uL with 100 uL chromatin aliquot. Incubate bead/antibody with chromatin using rotation for one hour at room temperature then move to 4*^◦^*C for another hour. After incubation, bead complexes were washed five times with a LiCl wash buffer and removed remaining ions with a single wash with 1 mL of cold TE buffer. Chromatin was eluted from beads by incubating with intermittent shaking for 1 hour at 65*^◦^*C in IP elution buffer, followed by incubating overnight at 65*^◦^*C to reverse formaldehyde cross-links. DNA was purified using DNeasy Blood and Tissue kit (Qiagen 69506) and eluted in a final volume of 50 uL EB. 2 uL was used to quantify recovered DNA using Qubit dsDNA HS Assay kit (Thermo Q32854). For input controls, one aliquot of each tissue was brought to 200 uL with RIPA and reverse-crosslinked overnight at 65*^◦^*C. The following morning, samples were incubated an additional 30 minutes with 20 uL Proteinase K and 4 uL RNase A (Qiagen 19101) and subsequently eluted for DNA using DNeasy Blood and Tissue kit.

The entirety of the remaining IP DNA (and 100 – 500 ng input control for sorted and bulk chromatin, respectively) were used to generate sequencing libraries. Libraries were prepared by blunting and ligating ChIP DNA fragments to Illumina sequencing adapters for amplification with barcoded primers (30 sec at 98*^◦^*C; [10 sec at 98*^◦^*C, 30 sec at 65*^◦^*C, 30 sec at 72*^◦^*C] x 15 cycles; 5 min at 72*^◦^*C). Libraries were quantified with Qubit dsDNA HS Assay kit and visualized with Standard Sensitivity NGS Fragment Analysis Kit (Advanced Analytical DNF-473) and Fragment Analyzer 5200 (Agilent). Libraries were sequenced using the Illumina NovaSeq with 100 bp single-end runs.

### ChIP-seq analysis pipeline

Prior to analysis, reads were processed to remove optical duplicates with clumpify (BBMap v38.20; Bushnell B.; https://sourceforge.net/projects/bbmap/) [dedupe=t optical=t dupedist=25 and remove adapter reads with Cutadapt^69^ (v1.16) [-a AGATCGGAAGAGC -m 40]. Input reads for TFs were capped at 30 million and 40 million for bulk and sorted, respectively, using Seqtk (v1.2; https://github.com/lh3/seqtk). Individual experiments were constructed following ENCODE guidelines70 with two donors used as biological replicates for each experiment unless otherwise noted. Results were analyzed with the ENCODE processing pipeline (https://github.com/ENCODE-DCC/chip-seq-pipeline2) with alignments to the hg38 build of the human genome. All software within the package was run using the default settings or those recommended by the authors for transcription factors [-type tf] or histones [-type histone]. Final peaks were called using the IDR näıve overlapping method with a threshold of 0.05. QC measurements and results of the reproducibility test provided by ENCODE are listed in Supplementary Table 2 with most experiments passing ENCODE recommended benchmarks. For completion, we include some experiments that failed in a certain cell-type if the tested TF passed in another experiment. Shortcomings in the reproducibility test were typically attributed to an experiment performing much better in one donor versus another.

### Whole genome bisulfite sequencing (WGBS)

DNA was extracted from 10-20 mg fresh-frozen postmortem brain tissue using the DNEasy Blood and Tissue kit (cat#: 69504, Qiagen) following the manufacturer’s instructions. WGBS dual indexed libraries were generated using NEBNext Ultra DNA library Prep kit for Illumina (New England BioLabs) according to the manufacturer’s instructions with modifications. 500 ng gDNA was quantified by Qubit dsDNA HS assay (Invitrogen) and 1% unmethylated lambda DNA (cat#: D1521, Promega) was spiked in to measure bisulfite conversion efficiency. DNA was fragmented to an average insert size of 400-450 bp using a Covaris S2 sonicator in a 55 uL volume. The fragmented gDNA was converted to end-repaired, adenylated DNA using the NEBNext Ultra End Repair/dA-Tailing Module (cat#: 7442L, New England BioLabs). Methylated adaptors (NEBNext Multiplex Oligos for Illumina; cat#: E7535L New England BioLabs) were ligated to the product from the preceding step using the NEBNext Ultra Ligation Module (cat#: 7445L, New England BioLabs). Size-selection was performed using AMPure XP beads and insert sizes of 400 bp were isolated (0.4x and 0.2x ratios). Samples were bisulfite converted after size selection using the EZ DNA Methylation-Lightning Kit (cat#: D5030, Zymo) following the manufacturer’s instructions. Amplification was performed following bisulfite conversion using primers from the NEBNext Multiplex Oligos for Illumina (cat#: E6440S, New England BioLabs) and the Kapa HiFi Uracil+ PCR system (cat#: KK2801, Kapa Biosystems) with the following cycling parameters: 98*^◦^*C 45 sec / 8 cycles: 98*^◦^*C 15 sec, 65*^◦^*C 30 sec, 72*^◦^*C 30 sec / 72*^◦^*C 1min. The PCR enriched product was cleaned up using 0.9x AMPure XP beads (cat#: A63881, Beckman Coulter). Final libraries were run on 2100 Bioanalyzer (Agilent) using the High-Sensitivity DNA assay; samples were also run on Bioanalyzer after shearing and size selection for quality control purposes. Libraries were quantified by qPCR using the Library Quantification Kit for Illumina sequencing platforms (cat#: KK4824, KAPA Biosystems, Boston), using 7900HT Real Time PCR System (Applied Biosystems). Libraries were sequenced with the Illumina NovaSeq S4 flowcell using 151 bp paired-end run with a 10% PhiX spike-in.

### Enzymatic Methyl-seq Kit of sorted nuclei

Nuclei were isolated from brain tissue samples and sorted similarly to sorted ChIP-seq experiments described above excluding the formaldehyde fixation step. DNA was extracted similarly to WGBS protocol above and processed using the NEBNext Enzymatic Methyl-seq Kit (NEB E7120S) according to the manufacturer’s recommendations. Libraries were sequenced with the Illumina NovaSeq S4 flowcell using 151 bp paired-end run with a 10% PhiX spike-in.

### Mapping and quality control of methylation reads

We trimmed reads of their adapter sequences using Trim Galore

(v0.6.0; http://www.bioinformatics.babraham.ac.uk/projects/trim galore/) and quality-trimmed using [– nextseq 20]. We then aligned these trimmed reads to the hg38 build of the human genome [including autosomes, sex chromosomes, mitochondrial sequence (available from https://software.broadinstitute.org/gatk/download/bundle) plus lambda phage (accession NC 001416.1) but excluding non-chromosomal sequences] using Bismark69 (v0.22.1) with the following alignment parameters: bismark –bowtie2 -X 1000 -1 *READ*1 *−* 2READ2.

We then used bismark methylation extractor to summarize the number of reads supporting a methylated cytosine and the number of reads supported a unmethylated cytosine for every cytosine in the reference genome. Specifically, we first computed and visually inspected the M-bias[5] of our libraries. Based on these results, we decided to ignore the first 5 bp of read2 in the subsequent call to bismark methylation extractor with parameters: –ignore r2 5 -p –bedGraph –counts –cytosine report –comprehensive –merge non CpG. The final cytosine report file summarizes the methylation evidence at each cytosine in the reference genome.

### Whole genome sequencing

High molecular weight was extracted from approximately 20mg of cortex tissue from each donor using either the MagAttract HMW DNA kit (Qiagen 67563) or Circulomics Nanobind Tissue kit (Pacbio 102-302-100). The linked read library was prepared using the Chromium Genome Reagent Kit v2 following the protocol provided by 10x Genomics. Sequence reads were processed using the longranger software suite from 10x Genomics. Linked reads were demultiplexed with longranger demux and then aligned to a 10x Genomics-provided, longranger-enabled hg38 reference (version 2.1.0) using longranger wgs v2.2.2. Variants were called using GATK 3.8-1-0-gf15c1c3ef via the –vcmode gatk option in the longranger wgs workflow.

### Assay for transposase-accessible chromatin using sequencing (ATAC-seq)

Homogenate, NeuN+, and NeuN-nuclei were isolated as previously described and 100,000 nuclei were used for ATAC-seq library preparation as per standard protocols^70–72^ with the following modifications. Briefly, 50 uL of cold 5X lysis buffer was added to sorted nuclei to reach a final concentration of 1X (10 mM Tris-HCl, pH 7.4, 10 mM NaCl, 3 mM MgCl2, 0.1% NP40) and incubated 20 min on ice followed by centrifugation for 10 min as previously described. The transposition reaction was incubated for 30 min or 1h at 37*^◦^*C (Illumina Tagment DNA Enzyme and Buffer kit; cat #:20034198). After column clean up via the Qiagen MinElute Reaction Cleanup kit (cat#:28204, Qiagen) and PCR amplification of libraries, an additional clean up with AMPure XP beads (0.8x ratio) was performed with two 80% ethanol washes before quantification using a DNA High Sensitivity chip on a 2100 BioAnalyzer (Agilent). Libraries were sequenced with the Illumina NovaSeq S2 50bp paired-end dual indexed run. Reads were processed using the standard ENCODE ATAC-seq pipeline (v1.7.0).

### RNA-seq

Total RNA was isolated using the Qiagen miRNeasy Mini kit according to the manufacturer’s protocol for acquiring total RNA with a DNAse digest, using approximately 20mg pulverized brain tissue or 200,000 nuclei as input. 10ng and 1ug of RNA (from nuclei and bulk, respectively) was used as input for TruSeq stranded Total RNA Library Prep Gold kit (Illumina 20020598) according to manufacturer’s protocol with TruSeq RNA UD Indexes (Illumina 20022371). Sequencing was carried out using Illumina NovaSeq S4 150bp pair-end dual indexed run. Reads were aligned to hg38-GENCODEv42 using STAR with ENCODE standard options [-outFilterType BySJout –outFilterMultimapNmax 20 –alignSJoverhangMin 8 –alignSJDBoverhangMin 1 –outFilterMismatchNmax 999 –outFilterMismatchNoverReadLmax 0.04 –alignIntronMin 20 –alignIntronMax 1000000 – alignMatesGapMax 1000000]. Count tables were generated using htseq-count [-m union -s no] then transformed to Transcripts Per Million (TPM).

### GREAT analysis

GREAT analysis was performed using the rGREAT (v2.1.8) package (https://www.bioconductor.org/packages/release/bioc/html/rGREAT.html). Genomic regions were associated with genes using the basal plus extension method (5kb upstream, 1kb downstream, 500kb max extension). Enrichment for GO Biological Process terms were calculated within GREAT with background regions set as the union of all ChIP-seq peaks.

### Correlation matrix and PCA analysis

The signal value for each TF at each peak was first quantified using the custom script *pull vals to GR.R* which calculates the median signal value from the fold-change bigWig within the specified peaks. PCA was performed on the matrix of signal values using the *prcomp* function with scaling. The correlation matrix was then calculated for all TFs using the first 20 PCs. Clusters were determined with ComplexHeatmap using k-means partitioning (k=3).

### Motif Calling

Motifs were identified with motifmatchr and a curated list of JASPAR 2022 motifs at a p-value cutoff of 5e-05. For motif centrality, peaks were centered on the summit and resized to 500 bp before calling motif positions. Motif distance was determined from the start of the motif to the summit of the peak. Centrality score was calculated as the variance in motif distance.

De novo motif calling was performed using the MEME suite (5.3.3) on the top 500 peaks (ranked by q-value from ENCODE pipeline) that were resized to 200 bp centered on the summit. MEME-ChIP was run with [-meme-nmotifs 5 -maxw 20 -minw 6 -meme-norand]. An experiment successfully called a motif de novo if the expected motif available in the JASPAR^47^ or CIS-BP^75^ database appears in the top 5 derived motif sequences.

For motif clustering, the MEME-ChIP suite was used to call de novo motifs using the top 500 peaks (sorted by q-value) and tested for enrichment and centrality in regions beyond those top peaks. The de novo motifs were then compared using TOMTOM and clustered based on similarity based on a previously published method^50^.

### Predictive Models

Predicted transcription factor binding scores were calculated using Virtual ChIP-seq *make input.p* and *predict.py*. ATAC-seq idr narrowPeak files and TPM transformed RNA-seq matrices were used as input for the predictions. Predictions were run for TFs that were both ChIPed in this study and had a pre-trained model available with Virtual ChIP-seq. Posterior probabilities were determined for a bin size of 200bp. Bins overlapping ENCODE blacklist regions were excluded (https://www.encodeproject.org/files/ENCFF419RSJ/@@download/ENCFF419RSJ.bed.gz). Precision and recall were calculated at a posterior probability step size of 0.01, and auPR was calculated using zoo’s *rollmean* function.

## References

1. Lambert, S. A. et al. The Human Transcription Factors. Cell 172, 650–665 (2018).

2. Partridge, E. C. et al. Occupancy maps of 208 chromatin-associated proteins in one human cell type. Nature 583, 720–728 (2020).

3. Lee, T. I. & Young, R. A. Transcriptional Regulation and Its Misregulation in Disease. Cell 152, 1237–1251 (2013).

4. Hammonds, A. S. et al. Spatial expression of transcription factors in Drosophilaembryonic organ development. Genome Biol. 14, R140 (2013).

5. Spitz, F. & Furlong, E. E. M. Transcription factors: from enhancer binding to developmental control. Nat. Rev. Genet. 13, 613–626 (2012).

6. Singh, H., Khan, A. A. & Dinner, A. R. Gene regulatory networks in the immune system. Trends Immunol. 35, 211–218 (2014).

7. Lee, R. van der Correard, S. & Wasserman, W. W. Deregulated Regulators: Disease-Causing cis Variants in Transcription Factor Genes. Trends Genet. 36, 523–539 (2020).

8. Carrasco Pro, S., Bulekova, K., Gregor, B., Labadorf, A. & Fuxman Bass, J. I. Prediction of genome-wide effects of single nucleotide variants on transcription factor binding. Sci. Rep. 10, 17632 (2020).

9. Singh, T. et al. Rare coding variants in 10 genes confer substantial risk for schizophrenia. Nature 604, 509–516 (2022).

10. Ramaker, R. C. et al. Post-mortem molecular profiling of three psychiatric disorders. Genome Med. 9, 72 (2017).

11. Szczepankiewicz, A. et al. Glucocorticoid receptor polymorphism is associated with major depression and predominance of depression in the course of bipolar disorder. J. Affect. Disord. 134, 138–144 (2011).

12. Forrest, M. P. et al. The Psychiatric Risk Gene Transcription Factor 4 (TCF4) Regulates Neurodevelopmental Pathways Associated With Schizophrenia, Autism, and Intellectual Disability. Schizophr. Bull. 44, 1100–1110 (2018).

13. Working, S. et al. Biological insights from 108 schizophrenia-associated genetic loci. Nature 511, 421–427 (2014).

14. Trubetskoy, V. et al. Mapping genomic loci implicates genes and synaptic biology in schizophrenia. Nature 604, 502–508 (2022).

15. Stefansson, H. et al. Common variants conferring risk of schizophrenia. Nature 460, 744–747 (2009).

16. Stahl, E. et al. Genomewide association study identifies 30 loci associated with bipolar disorder. bioRxiv 173062 (2018) doi:10.1101/173062.

17. Bryois, J. et al. Evaluation of chromatin accessibility in prefrontal cortex of individuals with schizophrenia. Nat. Commun. 9, 3121 (2018).

18. Fullard, J. F. et al. An atlas of chromatin accessibility in the adult human brain. Genome Res. 28, 1243–1252 (2018).

19. Gandal, M. J., Leppa, V., Won, H., Parikshak, N. N. & Geschwind, D. H. The road to precision psychiatry: translating genetics into disease mechanisms. Nat. Neurosci. 19, 1397–1407 (2016).

20. Cross-Disorder Group of the Psychiatric Genomics Consortium. Identification of risk loci with shared effects on five major psychiatric disorders: a genome-wide analysis. Lancet Lond. Engl. 381, 1371–1379 (2013).

21. PsychENCODE Consortium, et al. The PsychENCODE project. Nat. Neurosci. 18, 1707–1712 (2015).

22. Fromer, M. et al. Gene expression elucidates functional impact of polygenic risk for schizophrenia. Nat. Neurosci. 19, 1442–1453 (2016).

23. Girdhar, K. et al. Cell-specific histone modification maps in the human frontal lobe link schizophrenia risk to the neuronal epigenome. Nat. Neurosci. 21, 1126–1136 (2018).

24. Girdhar, K. et al. Chromatin domain alterations linked to 3D genome organization in a large cohort of schizophrenia and bipolar disorder brains. Nat. Neurosci. 25, 474–483 (2022).

25. The ENCODE Project Consortium, et al. Perspectives on ENCODE. Nature 583, 693–698 (2020).

26. Haenni, S. et al. Analysis of C. elegans intestinal gene expression and polyadenylation by fluorescence-activated nuclei sorting and 3’-end-seq. Nucleic Acids Res. 40, 6304–6318 (2012).

27. Dammer, E. B. et al. Neuron enriched nuclear proteome isolated from human brain. J. Proteome Res. 12, 3193–3206 (2013).

28. Dunham, I. et al. An integrated encyclopedia of DNA elements in the human genome. Nature 489, 57–74 (2012).

29. The ENCODE Project Consortium, et al. Expanded encyclopaedias of DNA elements in the human and mouse genomes. Nature 583, 699–710 (2020).

30. McLean, C. Y. et al. GREAT improves functional interpretation of cis-regulatory regions. Nat. Biotechnol. 28, 495–501 (2010).

31. Gu, Z. & Hübschmann, D. rGREAT: an R/bioconductor package for functional enrichment on genomic regions. Bioinformatics 39, btac745 (2023).

32. Datta, V., Siddharthan, R. & Krishna, S. Detection of cooperatively bound transcription factor pairs using ChIP-seq peak intensities and expectation maximization. PLoS ONE 13, e0199771 (2018).

33. Wei, B. et al. A protein activity assay to measure global transcription factor activity reveals determinants of chromatin accessibility. Nat. Biotechnol. 36, 521–529 (2018).

34. Dobson, T. H. W. et al. Regulation of USP37 Expression by REST-Associated G9a-Dependent Histone Methylation. Mol. Cancer Res. MCR 15, 1073–1084 (2017).

35. Mulligan, P. et al. CDYL Bridges REST and Histone Methyltransferases for Gene Repression and Suppression of Cellular Transformation. Mol. Cell 32, 718–726 (2008).

36. Hevner, R. F. et al. Tbr1 Regulates Differentiation of the Preplate and Layer 6. Neuron 29, 353–366 (2001).

37. Britanova, O. et al. Satb2 is a postmitotic determinant for upper-layer neuron specification in the neocortex. Neuron 57, 378–392 (2008).

38. Moffat, J. J. et al. Differential roles of ARID1B in excitatory and inhibitory neural progenitors in the developing cortex. Sci. Rep. 11, 3856 (2021).

39. Wiegreffe, C. et al. Bcl11a (Ctip1) Controls Migration of Cortical Projection Neurons through Regulation of Sema3c. Neuron 87, 311–325 (2015).

40. Hao, N., Bhakti, V. L. D., Peet, D. J. & Whitelaw, M. L. Reciprocal regulation of the basic helix-loop-helix/Per-Arnt-Sim partner proteins, Arnt and Arnt2, during neuronal differentiation. Nucleic Acids Res. 41, 5626–5638 (2013).

41. Latypova, X. et al. Haploinsufficiency of the Sin3/HDAC corepressor complex member SIN3B causes a syndromic intellectual disability/autism spectrum disorder. Am. J. Hum. Genet. 108, 929–941 (2021).

42. Fang, F. et al. A distinct isoform of ZNF207 controls self-renewal and pluripotency of human embryonic stem cells. Nat. Commun. 9, 4384 (2018).

43. Wreczycka, K. et al. HOT or not: examining the basis of high-occupancy target regions. Nucleic Acids Res. 47, 5735–5745 (2019).

44. Kvon, E. Z., Stampfel, G., Yáñez-Cuna, J. O., Dickson, B. J. & Stark, A. HOT regions function as patterned developmental enhancers and have a distinct cis-regulatory signature. Genes Dev. 26, 908–913 (2012).

45. Gerstein, M. B. et al. Integrative Analysis of the Caenorhabditis elegans Genome by the modENCODE Project. Science 330, 1775–1787 (2010).

46. Moorman, C. et al. Hotspots of transcription factor colocalization in the genome of Drosophila melanogaster. Proc. Natl. Acad. Sci. 103, 12027–12032 (2006).

47. Castro-Mondragon, J. A., et al. JASPAR 2022: the 9th release of the open-access database of transcription factor binding profiles. Nucleic Acids Res. 50, D165–D173 (2022).

48. Bailey, T. L., Johnson, J., Grant, C. E. & Noble, W. S. The MEME Suite. Nucleic Acids Res. 43, W39–W49 (2015).

49. McGann, J. C. et al. The Genome-Wide Binding Profile for Human RE1 Silencing Transcription Factor Unveils a Unique Genetic Circuitry in Hippocampus. J. Neurosci. 41, 6582–6595 (2021).

50. Vierstra, J. et al. Global reference mapping of human transcription factor footprints. Nature 583, 729–736 (2020).

51. Long, Y.-S. et al. Human transcription factor genes involved in neuronal development tend to have high GC content and CpG elements in the proximal promoter region. J. Genet. Genomics Yi Chuan Xue Bao 38, 157–163 (2011).

52. Mao, X., Yang, S.-H., Simpkins, J. W. & Barger, S. W. Glutamate receptor activation evokes calpain-mediated degradation of Sp3 and Sp4, the prominent Sp-family transcription factors in neurons. J. Neurochem. 100, 1300–1314 (2007).

53. Kiyama, T. et al. Essential roles of mitochondrial biogenesis regulator Nrf1 in retinal development and homeostasis. Mol. Neurodegener. 13, 56 (2018).

54. Worsley Hunt, R. & Wasserman, W. W. Non-targeted transcription factors motifs are a systemic component of ChIP-seq datasets. Genome Biol. 15, 412 (2014).

55. Dejosez, M. et al. Regulatory architecture of housekeeping genes is driven by promoter assemblies. Cell Rep. 42, (2023).

56. Karimzadeh, M. & Hoffman, M. M. Virtual ChIP-seq: predicting transcription factor binding by learning from the transcriptome. Genome Biol. 23, 126 (2022).

57. Barretina, J. et al. The Cancer Cell Line Encyclopedia enables predictive modelling of anticancer drug sensitivity. Nature 483, 603–607 (2012).

58. Anderson, A. G. et al. Single nucleus multiomics identifies ZEB1 and MAFB as candidate regulators of Alzheimer’s disease-specific cis-regulatory elements. Cell Genomics 0, (2023).

59. Finucane, H. K. et al. Partitioning heritability by functional annotation using genome-wide association summary statistics. Nat. Genet. 47, 1228–1235 (2015).

60. Managò, F. & Papaleo, F. Schizophrenia: What’s Arc Got to Do with It? Front. Behav. Neurosci. 11, 181 (2017).

61. Zhang, W. et al. Structural basis of arc binding to synaptic proteins: implications for cognitive disease. Neuron 86, 490–500 (2015).

62. Lanoue, V. et al. The adhesion-GPCR BAI3, a gene linked to psychiatric disorders, regulates dendrite morphogenesis in neurons. Mol. Psychiatry 18, 943–950 (2013).

63. Hass, J. et al. Associations between DNA methylation and schizophrenia-related intermediate phenotypes a gene set enrichment analysis. Prog. Neuropsychopharmacol. Biol. Psychiatry 59, 31–39 (2015).

64. Ohayon, S., Yitzhaky, A. & Hertzberg, L. Gene expression meta-analysis reveals the up-regulation of CREB1 and CREBBP in Brodmann Area 10 of patients with schizophrenia. Psychiatry Res. 292, 113311 (2020).

65. Liu, S. et al. The early growth response protein 1-miR-30a-5p-neurogenic differentiation factor 1 axis as a novel biomarker for schizophrenia diagnosis and treatment monitoring. Transl. Psychiatry 7, e998 (2017).

66. Savic, D., Gertz, J., Jain, P., Cooper, G. M. & Myers, R. M. Mapping genome-wide transcription factor binding sites in frozen tissues. Epigenetics Chromatin (2013) doi:10.1186/1756-8935-6-30.

67. Jiang, Y., Matevossian, A., Huang, H.-S., Straubhaar, J. & Akbarian, S. Isolation of neuronal chromatin from brain tissue. BMC Neurosci. 9, 42 (2008).

68. Reddy, T. E. et al. Genomic determination of the glucocorticoid response reveals unexpected mechanisms of gene regulation. Genome Res. 19, 2163–2171 (2009).

69. Martin, Marcel (Department of Computer Science, TU Dortmund, G. Cutadapt removes adapter sequences from high-throughput sequencing reads. EMBnet.journal 17, 10–12 (2011).

70. Landt, S. G. et al. ChIP-seq guidelines and practices of the ENCODE and modENCODE consortia. Genome Research (2012) doi:10.1101/gr.136184.111.

71. Krueger, F. & Andrews, S. R. Bismark: A flexible aligner and methylation caller for Bisulfite-Seq applications. Bioinformatics 27, 1571–1572 (2011).

72. Buenrostro, J. D., Giresi, P. G., Zaba, L. C., Chang, H. Y. & Greenleaf, W. J. Transposition of native chromatin for fast and sensitive epigenomic profiling of open chromatin, DNA-binding proteins and nucleosome position. Nat. Methods 10, 1213–1218 (2013).

73. Buenrostro, J. D., Wu, B., Chang, H. Y. & Greenleaf, W. J. ATAC-seq: A method for assaying chromatin accessibility genome-wide. Curr. Protoc. Mol. Biol. 2015, 21.29.1-21.29.9 (2015).

74. Corces, M. R. An improved ATAC-seq protocol reduces background and enables interrogation of frozen tissues. Nat. Methods 176, 139–148 (2017).

75. Weirauch, M. T. et al. Determination and inference of eukaryotic transcription factor sequence specificity. Cell 158, 1431–1443 (2014).

