## Supplemental Figures for "Extensive profiling of transcription factors in postmortem brains defines genomic occupancy in disease-relevant cell types and links TF activities to neuropsychiatric disorders"

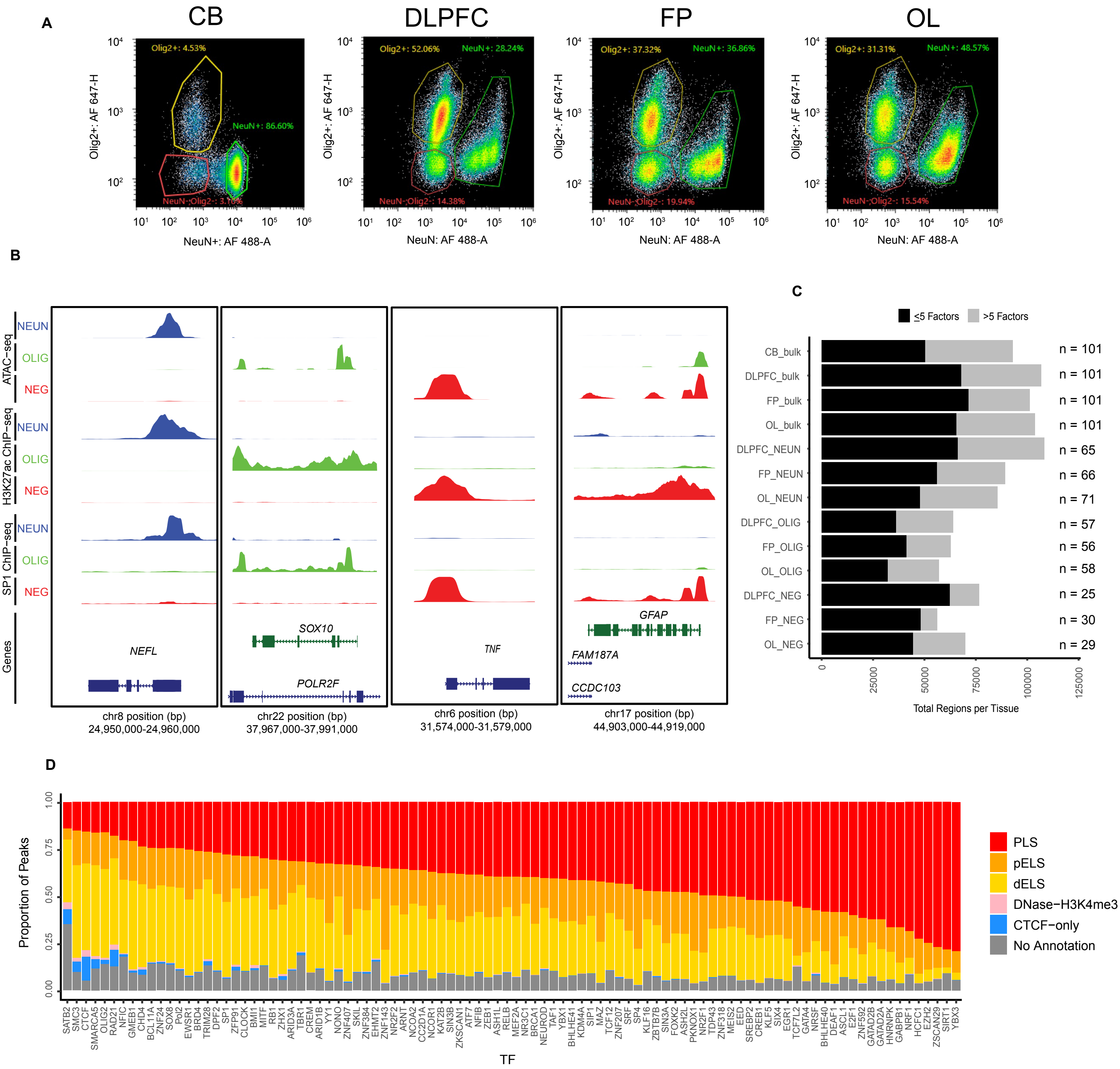


Figure S1. Supplemental to Figure 1. A) Example images of flow cytometry for nuclei sorted with NeuN and Olig2 antibodies with percentages of each sorted group. B) Browser tracks for ATAC-seq and ChIP-seq data over marker genes for neurons, oligodendrocytes, microglia, and astrocytes. C) Union of peaks summed for each tissue type from large brain regions with proportion bound by < or > 5 TFs. Total number of ChIP-seq experiments listed for each tissue cell-type. D) The proportion of peaks from each TF of DLPFC-bulk characterized by ENCODE cCREs.

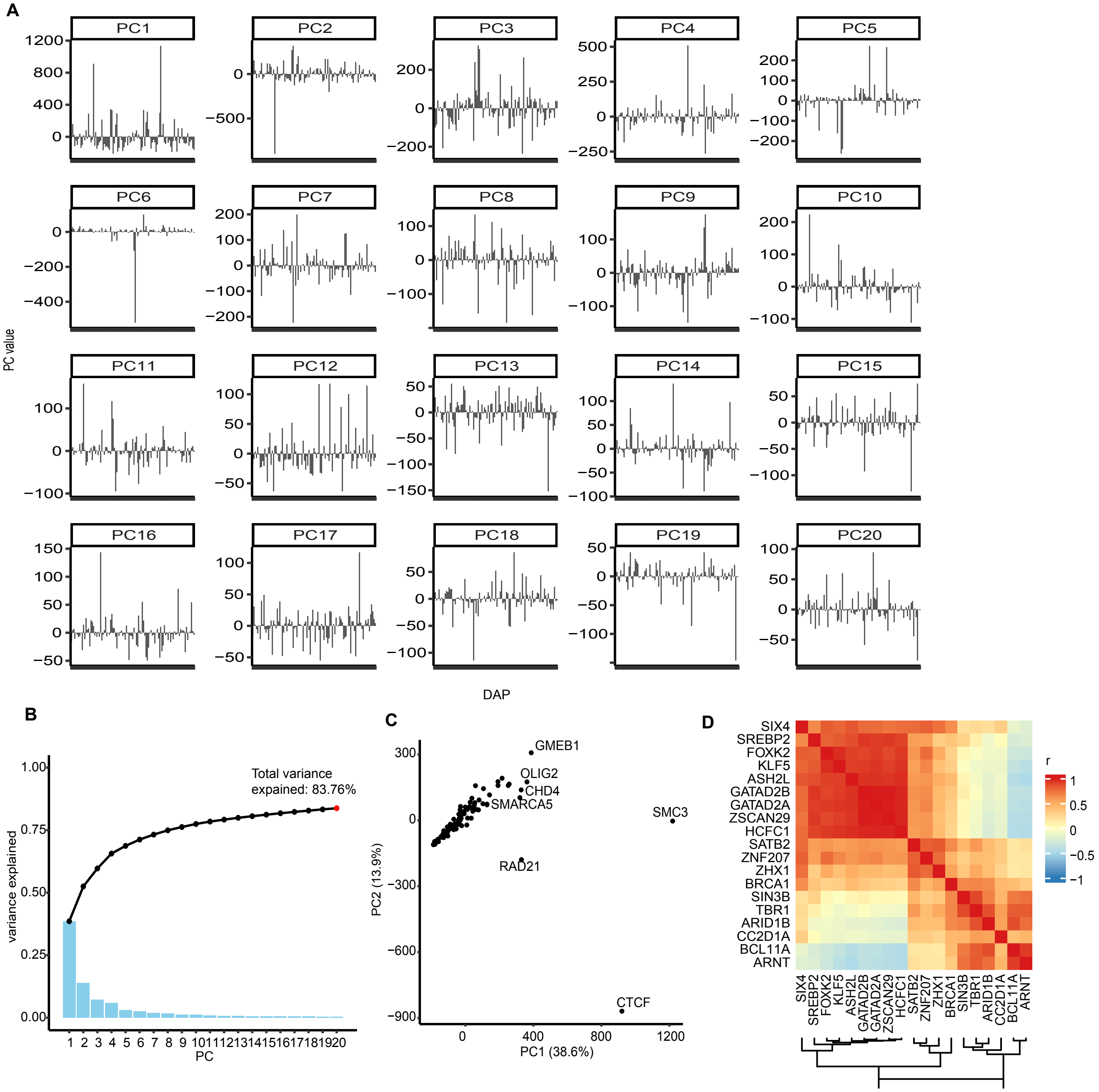


Figure S2. Supplemental to Figure 2. A) Values of first 20 PCs plotted for each TF from DLPFC-bulk. B) Visualization of variance from the first 20 PCs. Black line shows the cumulative variance explained. C) PC1 and PC2 of PCA from ChIP-seq of DLPFC-bulk. D) Group of correlated genes from main heatmap in Figure 2 highlighting genes related to neuronal regulation.

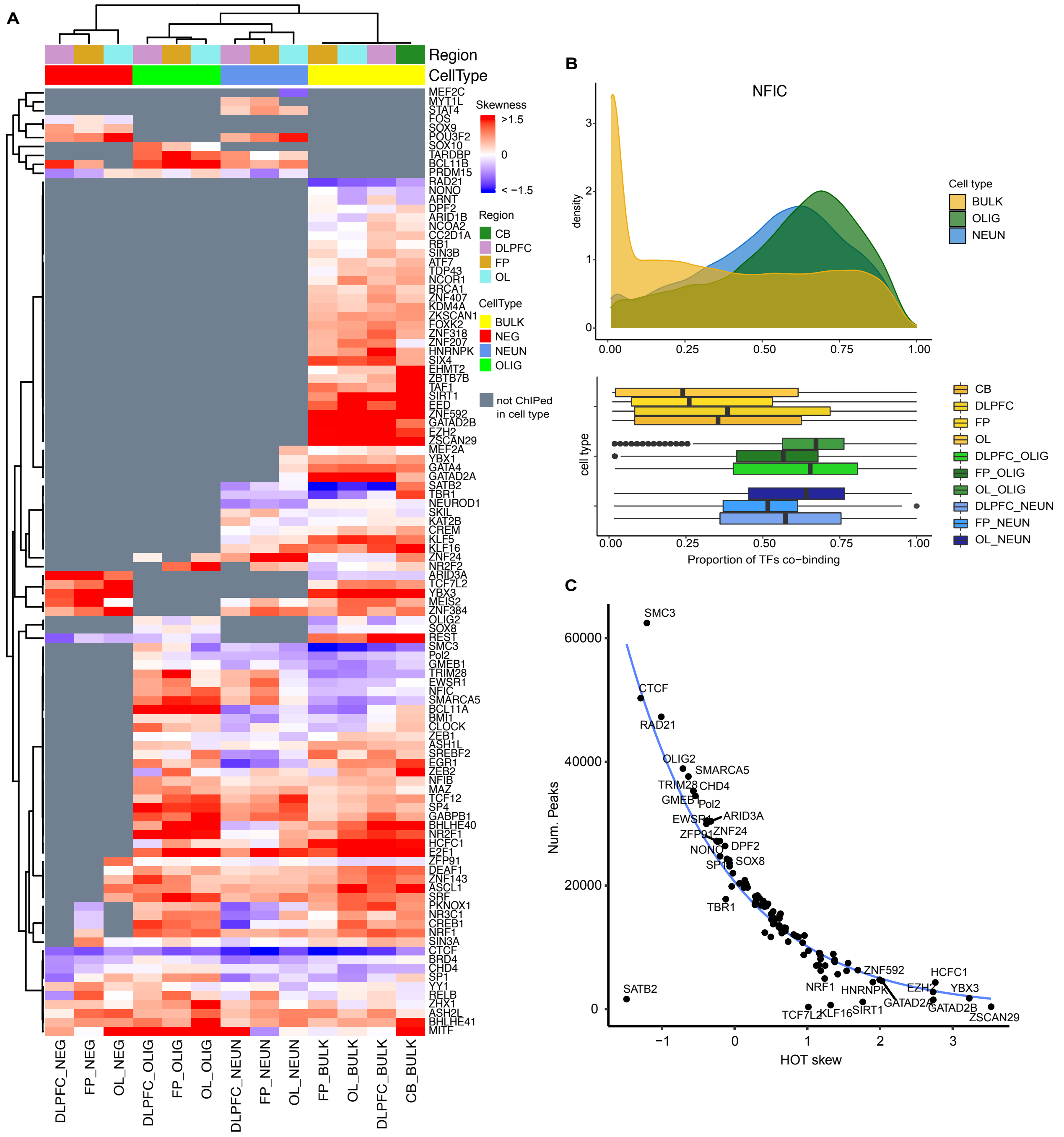


Figure S3. Supplemental to Figure 3. A) HOT skew values for ChIP-seq experiments across tissue and cell types clustered by skewness. B) Peak occupancy distribution for NFIC; average distribution (top) and individual distributions (bottom). C) Plot of HOT skew values versus the number peaks called from each ChIP-seq experiment in DLPFC-bulk.

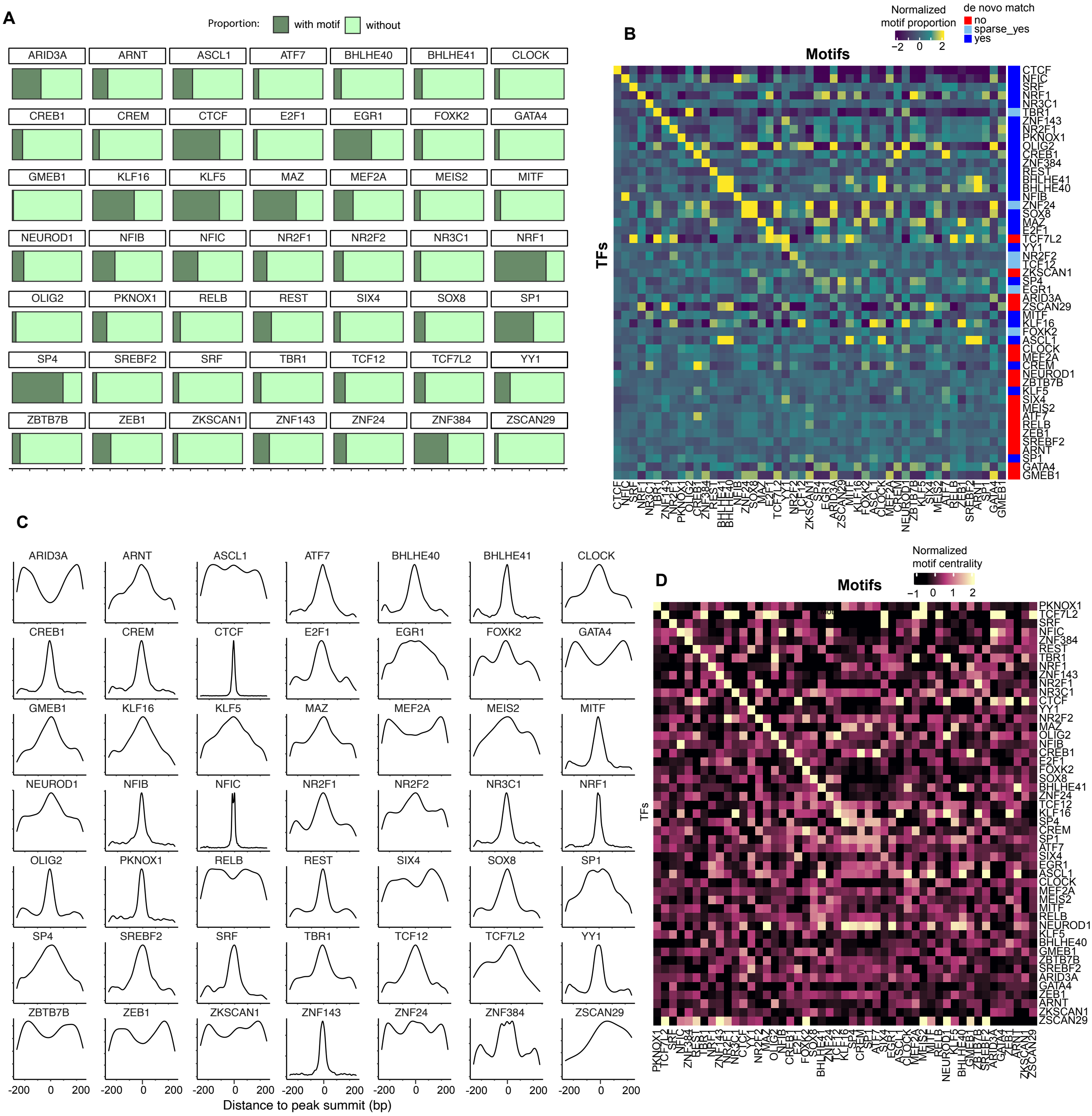


Figure S4. Supplemental to Figure 4. A) Proportion of peaks containing the expected motif from JASPAR 2022 database. B) Column-scaled proportion of peaks from each ChIP-seq matching all possible motifs tested. C) Centrality of expected motif calculated for each TF. D) Column-scaled centrality of motifs from each ChIP-seq matching all possible motifs tested.

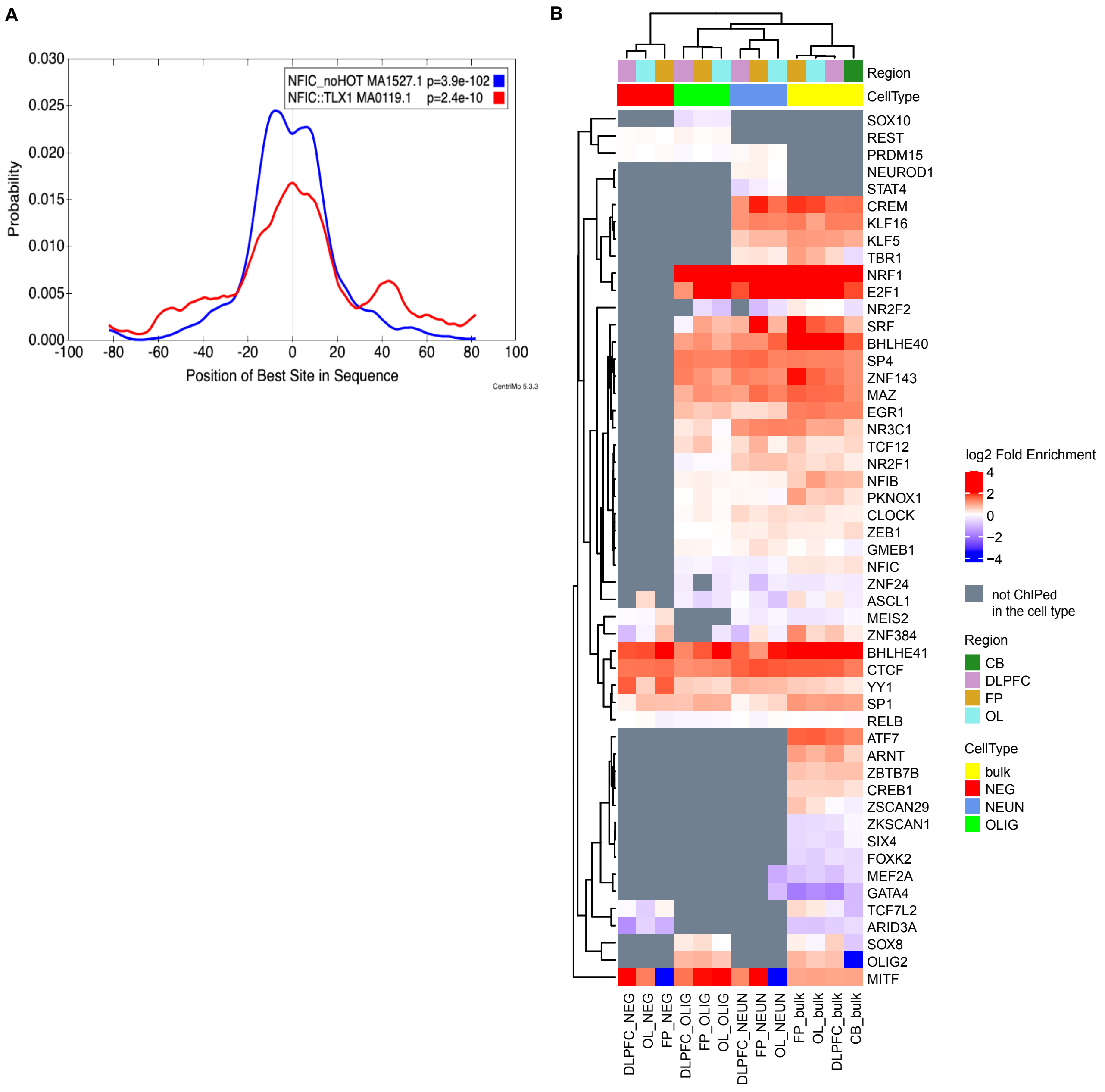


Figure S5. Supplemental to Figure 4 part 2. A) Centrality of NFIC before (red) and after (blue) removing HOT sites calculated using CENTRIMO with MEME-ChIP package. B) Enrichment of the expected motif in respective ChIP-seq peaks versus ATAC-seq peaks clustered by log2(Fold Enrichment).

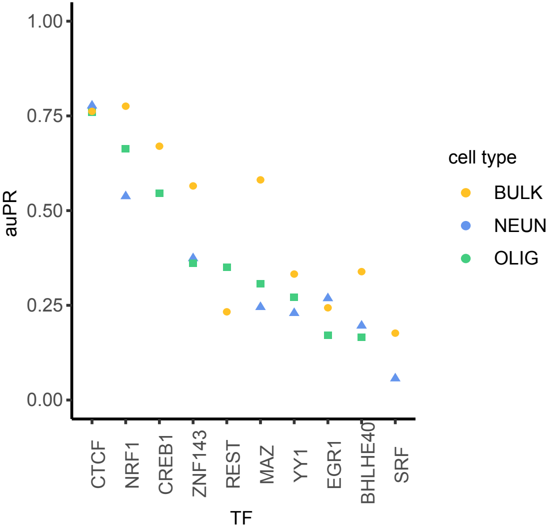


Figure S6. Supplemental to Figure 5. auPR of predictions for ChIP-seq experiments performed in bulk, NeuN+, and Olig2+ cell types from DLPFC.

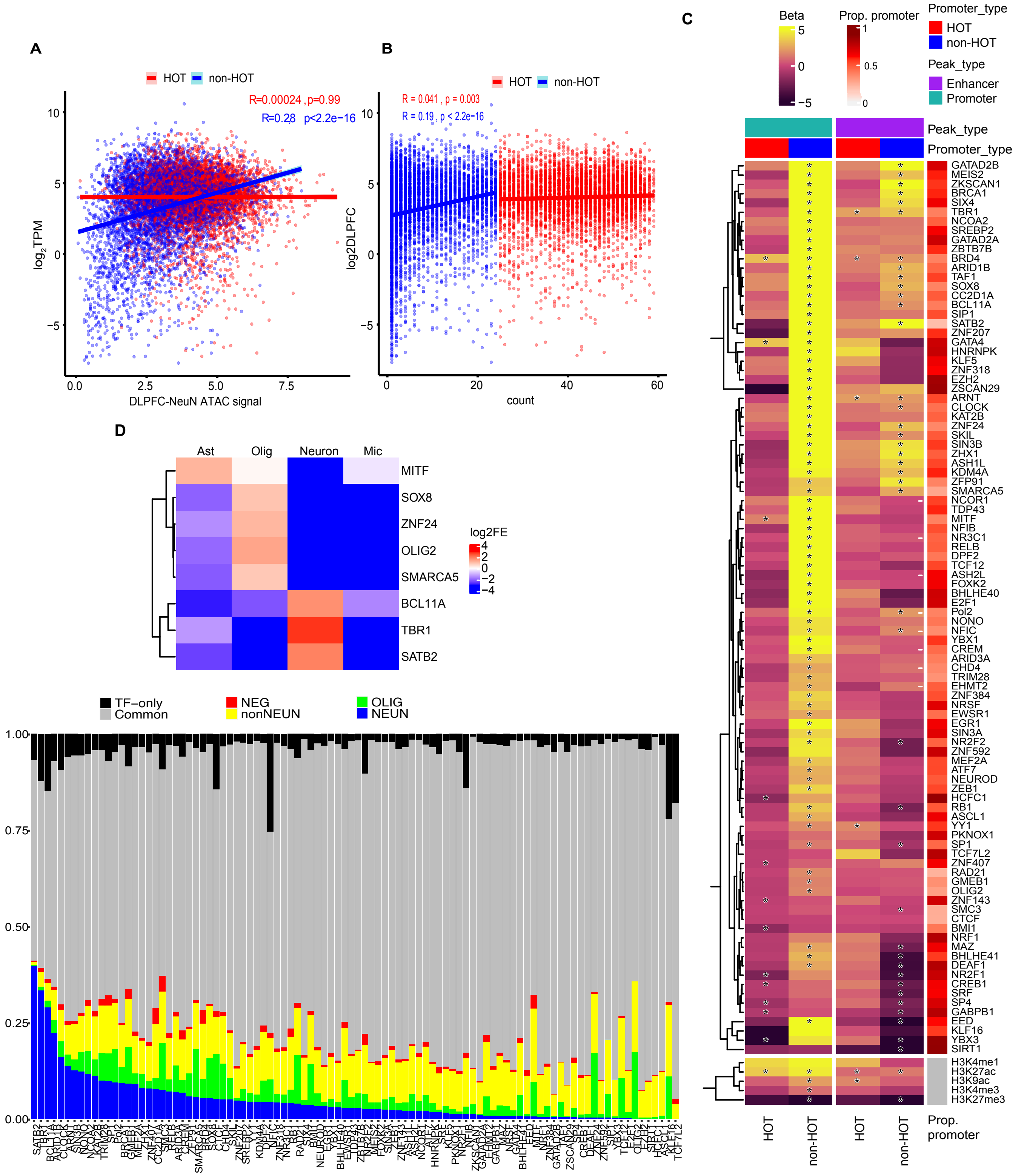


Figure S7. Supplemental to Figure 6. A) The correlation between gene expression in DLPFC-NEUN and ATAC-seq signal at promoters from NeuN+ nuclei segregated by whether the promoter overlaps a HOT site. B) Expression of each gene (log2 TPM) versus the number of TFs bound at the corresponding promoter. C) Heatmap similar to the main figure but with DLPFC-bulk data. D) Enrichment of ChIP-seq peaks of TFs for cell type-specific linked peaks identified in recent single-cell multiomic experiments from DLPFC. E) Similar to Figure 6D but with data from OL.

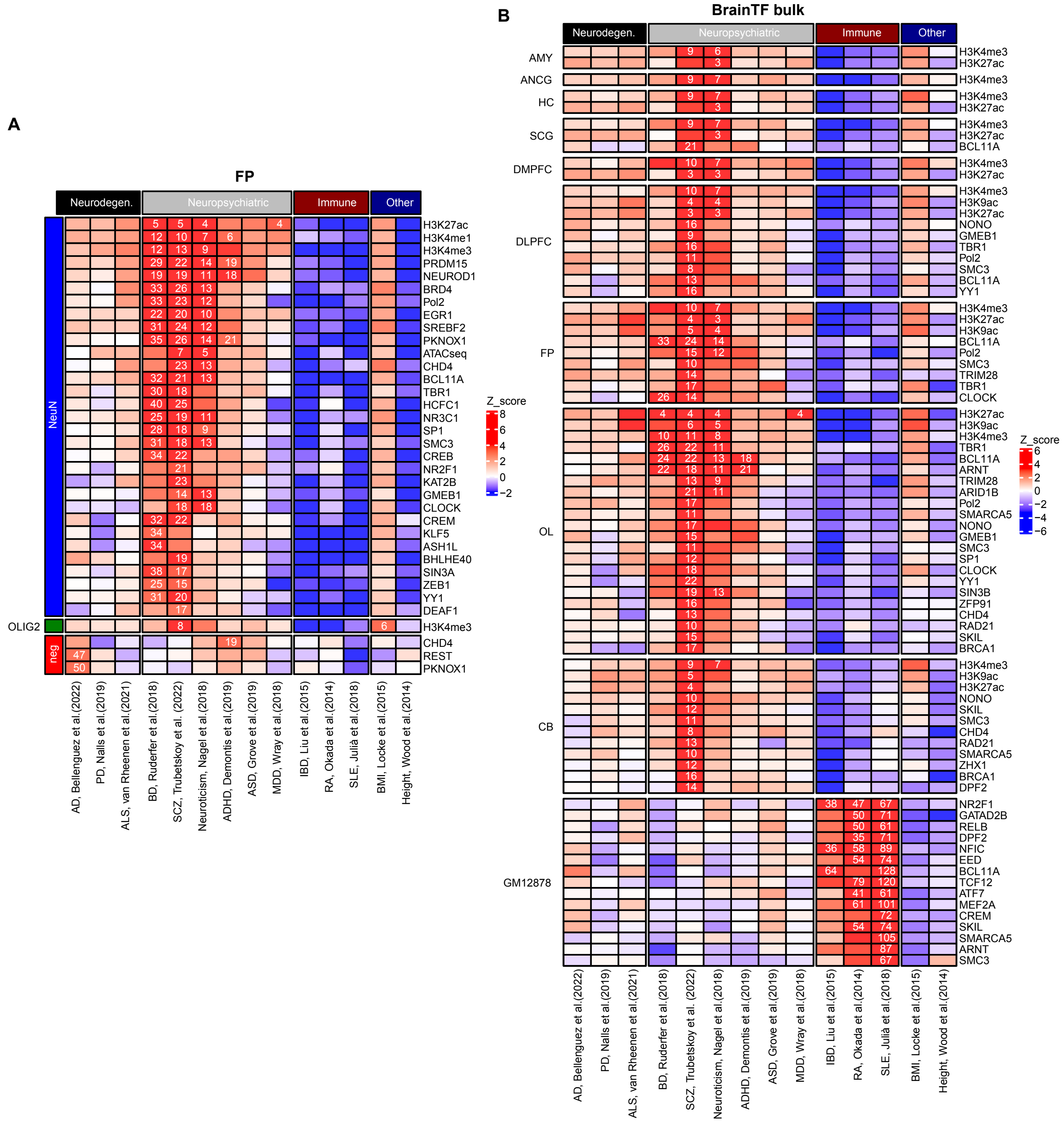


Figure S8. Supplemental to Figure 7. A) sLDSC results from FP. B) sLDSC results of all ChIP-seq data from bulk cell types alongside those from GM12878.
